# Intelliwaste: NMR of ^13^C-labeled Spent Media Enables Non-Invasive Metabolic Fingerprinting of Pluripotent Stem Cells and LIS1-Associated Neuropathology

**DOI:** 10.64898/2026.02.04.703261

**Authors:** Talia Harris, Maayan Karlinski Zur, Tamar Sapir, Orly Reiner, Rita Schmidt

## Abstract

Metabolic dysregulation is increasingly recognized as a key contributor to neurodevelopmental disorders. Here, we present ***Intelliwaste***, a non-invasive, cost-effective method for profiling carbon metabolism in pluripotent stem cells and brain organoids using ^13^C-labeled metabolites and ^1^H and ^13^C NMR spectroscopy. This approach enables longitudinal analysis of extracellular fluxes without disrupting cell viability.

We apply Intelliwaste to human embryonic stem cells (hESCs) cultured in a defined media enriched with >95% ^13^C_1_-Glucose. Under these conditions, ^13^C_3_-lactate emerged as the most abundant labeled product, with 20–50-fold lower fluxes to ^13^C_3_-alanine, ^13^C_2_-acetate, ^13^C_3_-serine, and ^13^C_3_-pyruvate, and 100–300-fold lower fluxes to ^13^C_1_-formate and multiple ^13^C-labeled glutamate species. These profiles allow for precise quantification of fractional metabolic isotopic labeling and glucose-derived carbon flow.

To demonstrate biological utility, we first examine the effect of ***L-glutamine omission***, which selectively reduces ^13^C_3_-alanine/^13^C_3_-lactate and ^13^C_4_-glutamate/^13^C_3_-lactate flux ratios, while the ^13^C_3_-Glutamate/^13^C_3_-Lactate and ^13^C_2_-Glutamate/^13^C_3_-Lactate flux ratios remained unchanged. These findings suggest a specific role for extracellular glutamine in modulating the activity of alanine aminotransferase and pyruvate carboxylase. We then characterized ***LIS1 mutant hESCs***—a model of lissencephaly—and observed significantly increased flux ratios involving ^13^C_4_-, ^13^C_3_-, and ^13^C_2_-glutamate relative to ^13^C_3_-lactate, indicating enhanced glutamate production via the TCA cycle.

These findings establish Intelliwaste as a powerful tool for metabolic profiling in the study of human neurodevelopment and disease. Its non-destructive nature makes it particularly well-suited for tracking metabolic changes during differentiation and in patient-derived organoid models of neurological disorders.

## Introduction

More than three decades ago, hemizygous deletion of the *Lissencephaly 1 (LIS1)* gene was identified as the genetic basis for lissencephaly in approximately 90% of patients with Miller-Dieker syndrome and 15% of patients with isolated lissencephaly ^1,2^. The neurodevelopmental consequences of *LIS1* haploinsufficiency have been extensively investigated in mouse models ^3^ and in cerebral organoids ^4,5^, revealing its critical role in multiple aspects of cortical development ^6^, particularly neuronal migration. More recent work has demonstrated that *LIS1* is also essential for stem cell viability: complete deletion leads to rapid cell death, whereas reduced LIS1 expression in heterozygous stem cells alters the expression of numerous genes, including those related to the extracellular matrix and pluripotency ^7^.

In this study, we sought to determine how decreased *LIS1* expression affects the metabolic phenotype of stem cells. This question arises from growing evidence that metabolic alterations contribute to neuropathologies observed in patient-derived pluripotent cells and cerebral organoids ^8–12^.

Traditionally, metabolic phenotypes are characterized by quantifying extracellular metabolite concentrations and by measuring intracellular RNA, protein, and metabolite levels in cell extracts^13–15^. However, neither approach alone provides a comprehensive picture: extracellular flux measurements are too coarse to reveal intracellular enzymatic activity, while intracellular concentration changes cannot be directly linked to metabolic fluxes. To address these limitations, the field of metabolic engineering has developed isotopic tracing methods, in which cells are incubated with isotopically labeled substrates; intracellular metabolites are then extracted, and their isotopic enrichment patterns are mapped onto biochemical pathways, enabling the estimation of fluxes through numerous intracellular enzymes^16–18^. Despite their power, these methods rely on destructive sampling of large cell masses – an endeavor that is prohibitively expensive for stem cell studies and unsuitable for longitudinal analyses of metabolic transitions during differentiation and organoid formation.

To overcome these challenges, we developed *Intelliwaste,* a non-destructive method for characterizing the metabolic phenotype of human embryonic stem cells (hESCs). In *Intelliwaste*, stem cells are routinely cultured in defined media containing specific ^13^C-enriched metabolite sites. Rather than discarding the spent media at the end of the incubation, it is collected and analyzed via ^1^H and ^13^C NMR spectroscopy to generate a metabolic fingerprint - hence the name *Intelliwaste*. Here, we demonstrate how *Intelliwaste*, using media enriched with ^13^C_1_-Glucose (^13^C_1_-Glc), can be used to characterize the metabolic phenotype of hESCs and to examine the impact of the LIS1 heterozygous deletion on this phenotype. Because *Intelliwaste* enables longitudinal analysis of extracellular fluxes without compromising cell viability, it can be readily extended to studies of differentiated stem cells and stem cell-derived organoids.

## Results

The experimental workflow involved standardized cell culture protocols, optimized procedures for robust spent media collection, high-resolution ^1^H and ^13^C NMR spectroscopy, and a dedicated pipeline for data processing and analysis (Figure 1). The Methods section provides detailed descriptions of each step.

**Figure 1:**
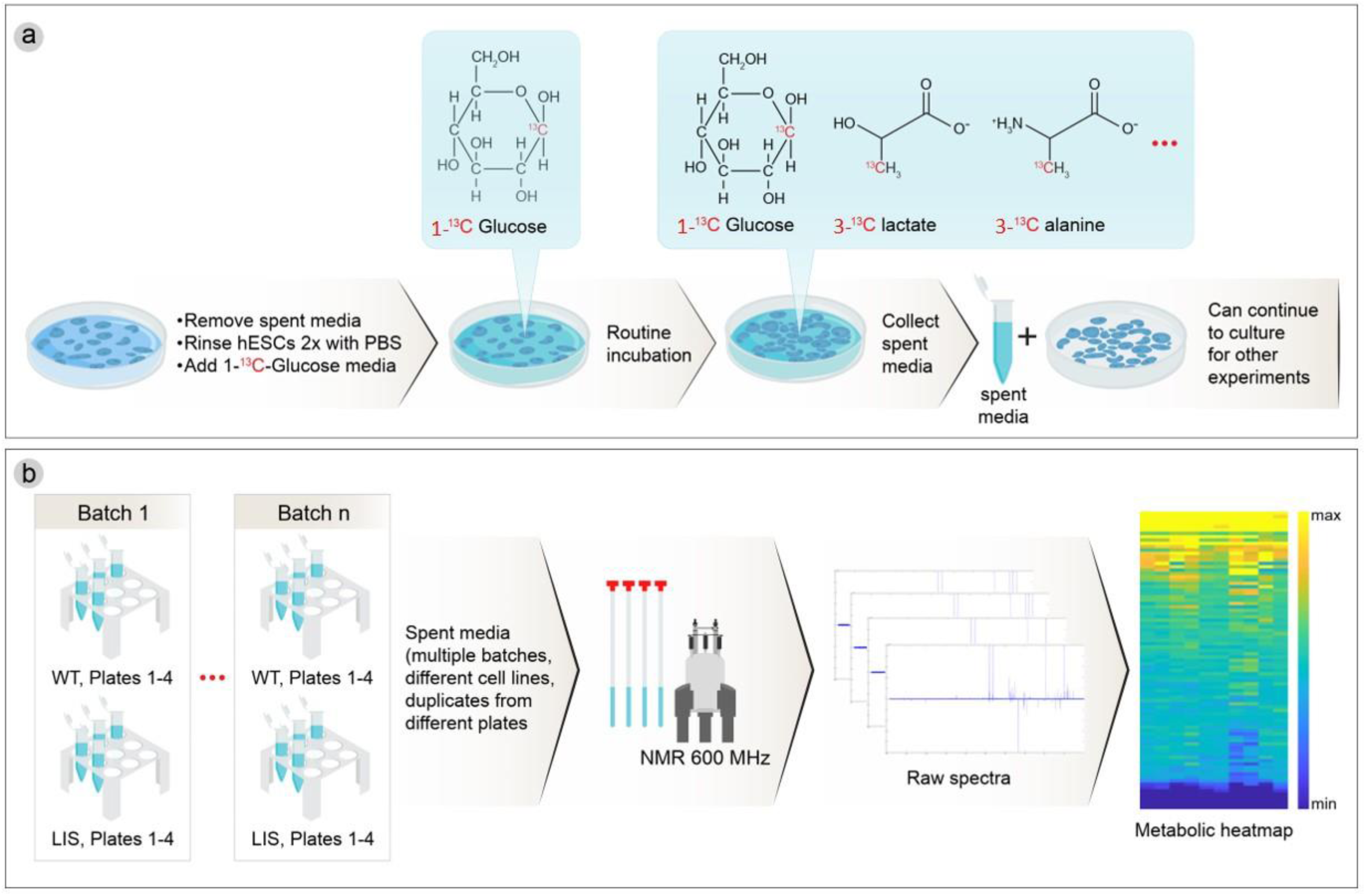
Schematic overview of the Intelliwaste workflow. (A) cell culture workflow showing the steps of isotopic labeling, routine incubation and spent media collection. (B) NMR spectroscopy acquisition and data processing pipeline, preparing multiple batches, collecting ^1^H and ^13^C NMR spectra, analyzing the peaks integrals and generating metabolic heatmaps.

### Metabolic Fingerprint of WT hESCs in Standard Media

Using our pH robust ^13^C NMR fitting pipeline, we successfully quantified 97 independent ^13^C metabolite sites from ^13^C NMR spectra of media incubated with and without hESCs (See Supplementary Table S1). Figure 2A shows a representative region of a raw spectrum overlaid with the corresponding Lorentzian fit, with several metabolite sites of interest labeled. Because spectra were acquired using a DEPT sequence, ^13^**C**H_3_ and ^13^**C**H sites appear as positive peaks (Figure 2A, blue labels) and ^13^**C**H_2_ and ^13^**C** sites appear as negative peaks (Figure 2A, red labels).

**Figure 2:**
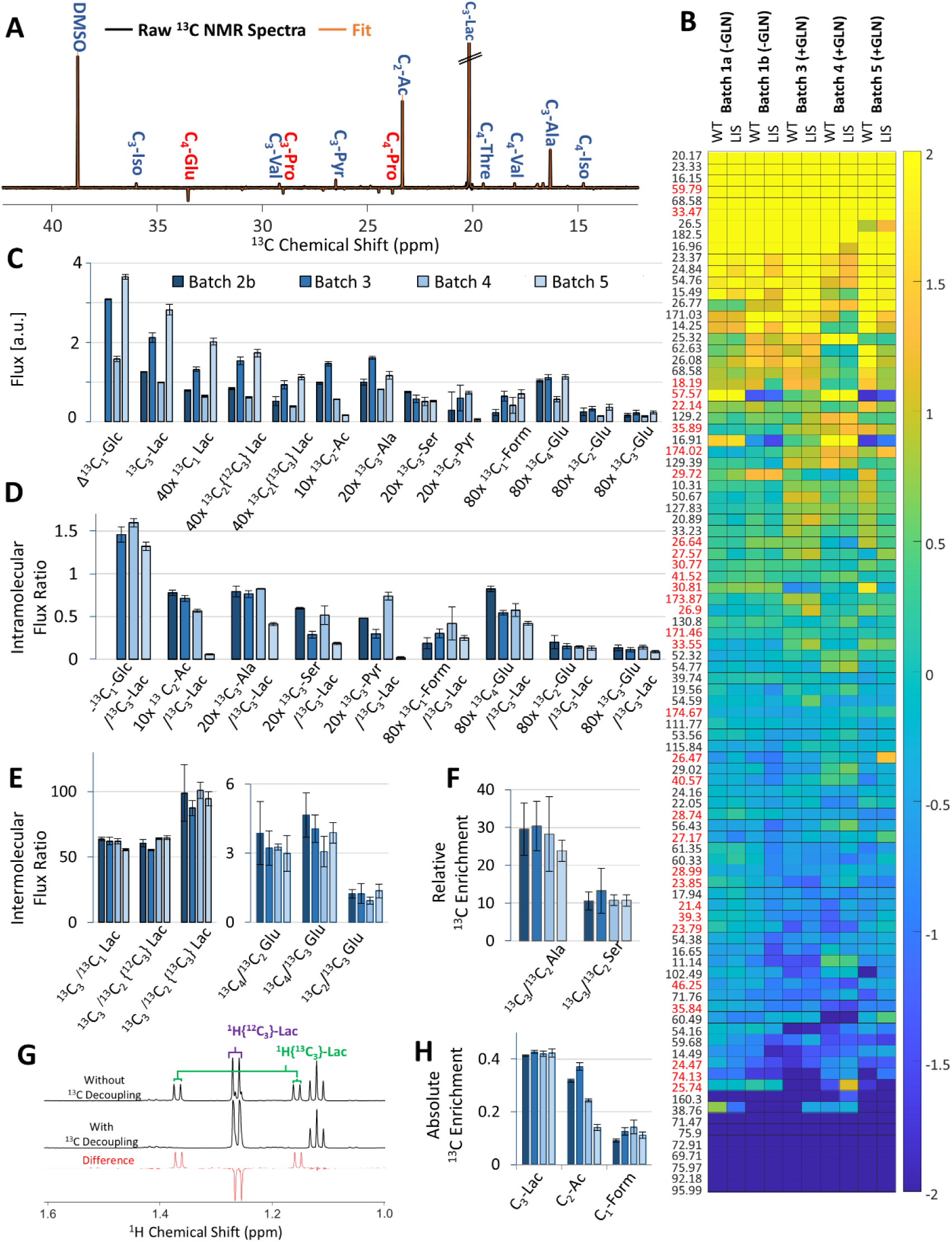
Overview of metabolic information extracted from ^13^C NMR measurements. (A) Representative segment of a ^13^C NMR spectrum (black) with corresponding automatic fit (orange). Peaks are positive for **C**H and **C**H_3_ (blue labels) and negative for C and CH2 (red labels). (B) The heat map shows changes in concentration for all 97 fitted metabolite sites in media incubated with hESCs, relative to media incubated without hESCs. Peaks are ordered by average change in signal intensity; yellow indicates the largest increases and blue indicates the largest decreases. (C) ^13^C Metabolic fluxes for 13 metabolite peaks that exhibited significant changes upon incubation with hESCs. Glucose peaks showed negative fluxes (consumption), while all others showed positive fluxes (production). (D) Intermolecular Metabolic Flux Ratios. (E) Intramolecular ^13^C Metabolic Flux ratios for Lactate (left) and Glutamate (right). (F) Relative ^13^C enrichment of C_3_ vs C_2_ positions for Alanine and Serine. (G) Example of ^1^H NMR spectra acquired without (top) and with (middle) ^13^C decoupling. The difference spectrum (bottom) reveals satellite peaks as negative signals flanking the central ^1^H{^12^C} peak. (H) Absolute enrichment calculated from satellite-to-total peak intensity ratio for C_1_-β-Glc, C_3_-Lac, C_2_-Ac, and C_1_-Form.

Based on differences in peak integrals between media incubated with and without hESCs (Figure 2B), 12 peaks exhibited positive metabolic fluxes (i.e. products), while 11 peaks exhibited negative metabolic fluxes (i.e. substrates). Among the twelve product peaks, four were assigned to Lactate (Lac) (^13^C_1_-Lac, ^13^C_2_-[^12^C_3_]-Lac, ^13^C_2_-[^13^C_3_]-Lac, ^13^C_3_-Lac); three corresponded to Glutamate (Glu) (^13^C_2_-Glu, ^13^C_3_-Glu and ^13^C_4_-Glu); the remaining product peaks were identified as ^13^C_2_-Acetate (Ac), ^13^C_3_-Alanine (Ala), ^13^C_3_-Serine (Ser), ^13^C_3_-Pyruvate (Pyr) and ^13^C_1_-Formate (Form).

The 11 peaks (that exhibited negative metabolic fluxes) corresponded to various positions in glucose (Glc), including the isotopically enriched ^13^C_1_ position and the natural abundance C2–C6 positions from the two Glc anomers. The range of observed metabolic ^13^C-labeled fluxes in the media spanned three orders of magnitude, with ^13^C_1_-Glc consumption and ^13^C_3_-Lac production representing the highest fluxes. The other quantified fluxes were 1-3 orders of magnitude smaller (Figure 2C, data shown for four representative experimental batches; full dataset is added in Supplementary Figures S2-5).

We observed good agreement across replicates from different plates of the same batch, as reflected in the small standard deviations in Figure 2C. Note: For Batch 2b, the ^13^C_1_-Glucose flux was not quantified, as its control media was inadvertently discarded. In contrast to other metabolites with similar initial concentrations, glucose showed higher variability across media preparations.

Flux measurements varied by more than two-fold across experimental batches, likely reflecting differences in initial cell number and proliferation rates during the 24-hour incubation period (Figure 2C). To account for this variability and establish a robust, cell-number-independent, metabolic fingerprint, we defined a series of intermolecular flux ratios (Figure 2D) and intramolecular flux ratios (Figure 2E). For serine (Ser) and alanine (Ala), intramolecular flux ratios could not be determined due to detected flux only in the C3 position; however, the relative ^13^C enrichment was estimated comparing the integrals of the C_2_ and C_3_ positions (Figure 2F).

In ^1^H NMR spectra, the peak of a proton bound to ^13^C is split into two peaks, separated from the central peak by 100-150 Hz, while ^13^C decoupling removes this splitting. Using the ratio of the ^1^H{^12^C} to ^1^H{^13^C} peaks, the absolute ^13^C enrichment of C_1_-β-Glc, C_3_-Lac, C_2_-Ac and C_1_-Form was determined, as demonstrated for C_3_-Lac in Figure 2G. ^13^C enrichment of C_1_-β-Glc consistently exceeded 95% across all batches (Supplementary Figure S5), confirming successful replacement of unlabeled spent media with freshly prepared ^13^C_1_-Glc labeled media. The observed ^13^C fractions for C_3_-Lac, C_2_-Ac and C_1_-Form across batches are shown in Figure 2H, with full dataset in Supplementary Figure S5.

Aggregating data from multiple independent experiments, we defined a representative metabolic fingerprint for WT hESCs cultured in standard media (Figure 3 and Table 1). In brief, consumption of 1460 ± 140 units of ^13^C_1_-Glc resulted in the production of:

- 1000 units of ^13^C_3_-Lac,
- 53 ± 33 units of ^13^C_2_-Ac,
- 35 ± 10 units of ^13^C_3_-Ala,
- 20 ± 10 molecules of ^13^C_3_-Ser,
- 19 ± 5 units of ^13^C_3_-Pyr,
- 3.6 ± 1.2 units of ^13^C_1_-Form,
- 7.4 ± 2.1 units of ^13^C_4_-Glu,
- 2.0 ± 0.4 units of ^13^C_2_-Glu
- 1.5 ± 0.3 units of ^13^C_3_-Glu.

**Figure 3:**
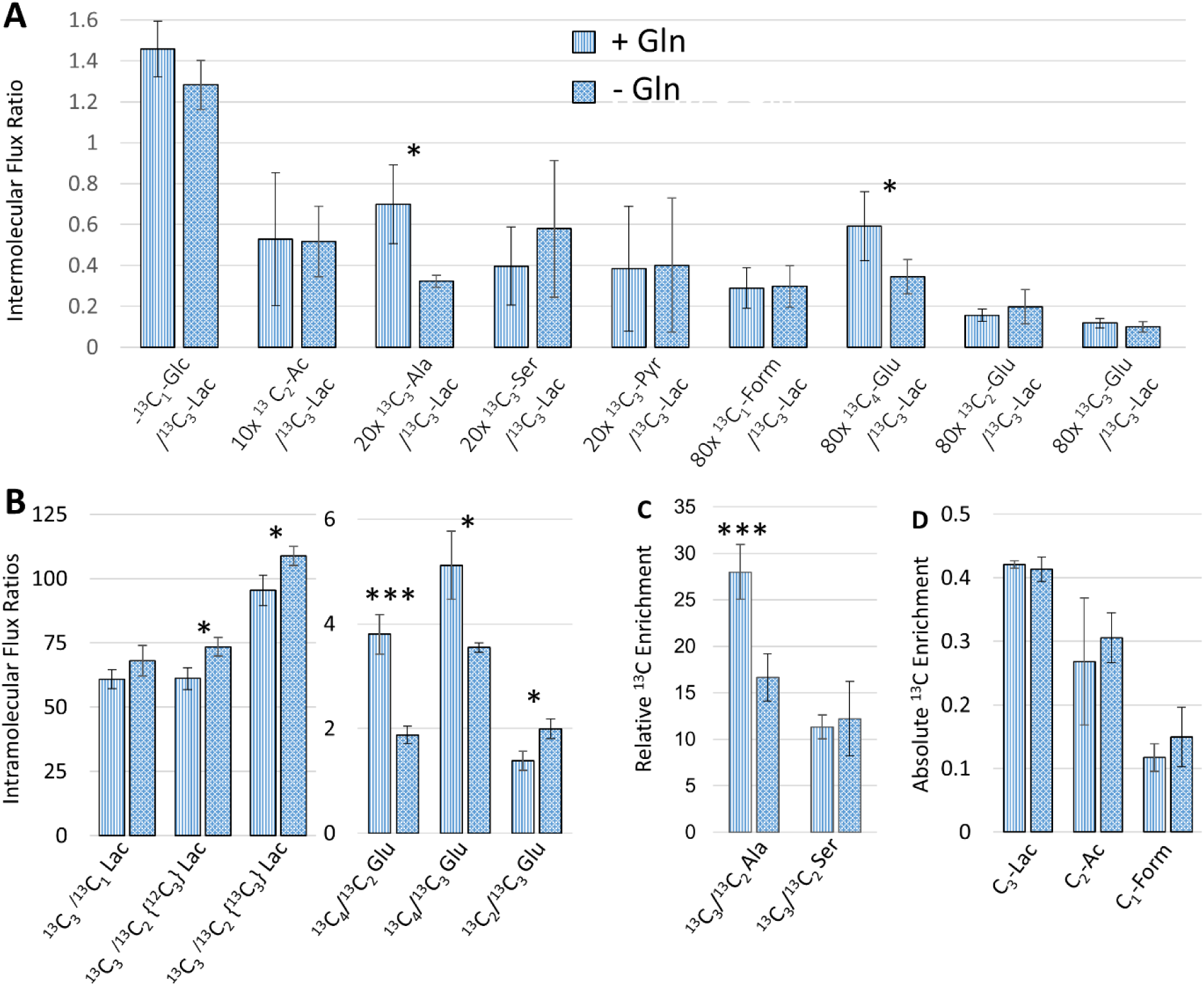
Metabolic fingerprint of WT hESCs cultured in standard media with L-Glutamine (n=4) and without L-Glutamine (n=3). (A) Intermolecular flux ratios, (B) intramolecular flux ratios (C) relative ^13^C enhancement, determined from the ratio of signal intensities between isotopically enriched and natural abundance positions. (D) absolute ^13^C enrichment, calculated from ^1^H-^13^C satellite peaks intensities in ^1^H NMR spectra. * p<0.1; ** p<0.01, ***p<0.005.

**Table 1:** Metabolic flux ratios and 13C Enrichment for WT hESCs cultured in standard media, with L-Glutamine (n=4) and without L-Glutamine (n=3)

|  |  |  |  |  |  |  |
| --- | --- | --- | --- | --- | --- | --- |
|  |  | + L-Glut |  | - L-Glut |  | p value * |
| Intermolecular Metabolic Flux Ratio | $^{13}\text{C}_1\text{-Glc}/^{13}\text{C}_3\text{-Lac}$ | 1.46 | ± 0.14 | 1.28 | ± 0.12 | |
| | $^{13}\text{C}_2\text{-Ac}/^{13}\text{C}_3\text{-Lac}$ | 0.053 | ± 0.033 | 0.052 | ± 0.017 | |
| | $^{13}\text{C}_3\text{-Ala}/^{13}\text{C}_3\text{-Lac}$ | 0.035 | ± 0.010 | 0.016 | ± 0.001 | p=0.02 |
| | $^{13}\text{C}_3\text{-Ser}/^{13}\text{C}_3\text{-Lac}$ | 0.020 | ± 0.010 | 0.029 | ± 0.017 | |
| | $^{13}\text{C}_3\text{-Pyr}/^{13}\text{C}_3\text{-Lac}$ | 0.019 | ± 0.015 | 0.020 | ± 0.0164 | |
| | $^{13}\text{C}_3\text{-Form}/^{13}\text{C}_3\text{-Lac}$ | 0.0036 | ± 0.0012 | 0.0037 | ± 0.0013 | |
| | $^{13}\text{C}_4\text{-Glu}/^{13}\text{C}_3\text{-Lac}$ | 0.0074 | ± 0.0021 | 0.0043 | ± 0.0010 | p=0.07 |
| | $^{13}\text{C}_2\text{-Glu}/^{13}\text{C}_3\text{-Lac}$ | 0.0020 | ± 0.0004 | 0.0025 | ± 0.0011 | |
| | $^{13}\text{C}_3\text{-Glu}/^{13}\text{C}_3\text{-Lac}$ | 0.0015 | ± 0.0003 | 0.0012 | ± 0.0003 | |
| Intramolecular Metabolic Flux Ratios | $^{13}\text{C}_3/^{13}\text{C}_1\text{-Lac}$ | 60.9 | ± 3.7 | 65.3 | ± 4.8 | |
| | $^{13}\text{C}_3/^{13}\text{C}_2\{^{12}\text{C}_3\}\text{-Lac}$ | 61.1 | ± 4.3 | 66.7 | ± 8.2 | p=0.01 |
| | $^{13}\text{C}_3/^{13}\text{C}_2\{^{13}\text{C}_3\}\text{-Lac}$ | 95.4 | ± 6.0 | 102.3 | ± 8.8 | p=0.02 |
| | $^{13}\text{C}_4/^{13}\text{C}_2\text{-Glu}$ | 3.8 | ± 0.6 | 1.9 | ± 0.2 | p=0.003 |
| | $^{13}\text{C}_4/^{13}\text{C}_3\text{-Glu}$ | 5.1 | ± 0.9 | 3.5 | ± 0.3 | p=0.04 |
| | $^{13}\text{C}_2/^{13}\text{C}_3\text{-Glu}$ | 1.4 | ± 0.2 | 2.0 | ± 0.4 | p=0.05 |
| Relative $^{13}\text{C}$ Enrichment | $^{13}\text{C}_3/^{13}\text{C}_2\text{-Ala}$ | 28 | ± 3 | 17 | ± 3 | p=0.003 |
| | $^{13}\text{C}_3/^{13}\text{C}_2\text{-Ser}$ | 11 | ± 1 | 12 | ± 4 | |
| Absolute $^{13}\text{C}$ Enrichment | $\text{C}_1\text{-Glc}$ | 0.98 | ± 0.01 | 0.98 | ± 0.01 | |
| | $\text{C}_3\text{-Lac}$ | 0.42 | ± 0.01 | 0.41 | ± 0.02 | |
| | $\text{C}_2\text{-Ac}$ | 0.27 | ± 0.10 | 0.31 | ± 0.04 | |
| | $\text{C}_1\text{-Form}$ | 0.12 | ± 0.02 | 0.15 | ± 0.05 | |
| *only p<0.1 are shown for simplicity |  |  |  |  |  |  |

This corresponds to intramolecular flux ratios of:

- 3.8 ± 0.6 for ^13^C_4_/^13^C_2_-Glu,
- 5.1 ± 0.9 for ^13^C_4_/^13^C_3_ Glu
- 1.4 ± 0.2 for ^13^C_2_/^13^C_3_ Glu.

In addition, the intramolecular flux ratios were:

- ^13^C_3_/^13^C_1_-Lac: 60.9 ± 3.7
- ^13^C_3_/^13^C_2_-{^12^C_3_}-Lac: 61.1 ± 4.3
- ^13^C_3_/^13^C_2_-{^13^C_3_}-Lac: 95.4 ± 6.0.

Finally, the observed ^13^C fractions were:

- C_3_-Lac: 0.42 ± 0.01,
- C_2_-Ac: 0.27 ± 0.10
- C_1_-Form: 0.12 ± 0.02.

Relative enrichment revealed that C_3_-Ala was 28 ± 3 fold more ^13^C enriched than C_2_-Ala, and C_3_-Ser was 11 ± 1 fold more ^13^C enriched than C_2_-Ser.

### Effect of Omitting L-Glutamine on Metabolic Fingerprint of WT hESCs

To increase the dimensionality of the WT hESC metabolic phenotype, we also characterized a metabolic fingerprint under conditions of L-glutamine (L-Gln) omission. Cells were incubated for 24 hours in ^13^C_1_-Glc enriched media with (+Gln) and without (-Gln) supplementation. The resulting metabolic fingerprints are summarized in Table 1 and Figure 3, highlighting significant differences (p ≤ 0.1). L-Gln omission resulted in a significant reduction in two key intermolecular flux ratios - ^13^C_3_-Ala/^13^C_3_-Lac (p=0.02) and ^13^C_4_-Glu/^13^C_3_-Lac (p=0.07).

Additionally, several intramolecular flux ratios were altered, observing:

- Increased flux ratios of ^13^C_3_/^13^C_2_-[^12^C_3_]-Lac (p=0.01), ^13^C_3_/^13^C_2_-[^13^C_3_]-Lactate (p=0.02), ^13^C_2_/^13^C_3_-Glu (p=0.05)
- Decreased flux ratios of ^13^C_4_/^13^C_2_-Glu (p=0.003) and ^13^C_4_/^13^C_3_-Glu (p=0.04).

Finally, glutamine omission also significantly reduced the relative enhancement of ^13^C_3_/^13^C_2_-Ala (p=0.003).

Together, these findings demonstrate that L-glutamine availability significantly influences carbon flux distribution, specifically decreasing the relative activities of alanine aminotransferase and pyruvate dehydrogenase, underscoring the metabolic adaptability of naïve human embryonic stem cells.

### Effect of *LIS1* mutation on the Metabolic Phenotype of hESCs

We next applied the Intelliwaste method to investigate how the *LIS1*mutation (clone 10F) affects the metabolic phenotype of naïve hESCs. This mutation has previously been shown to cause a 69 ± 12 % reduction in LIS1 protein expression, and cortical brain organoids derived from these hESCS recapitulate the hallmark smooth brain phenotype characteristic of lissencephaly ^4^.

In hESCs cultured in standard media, the LIS1 mutation led to a significant increase in several key metabolic flux ratios when compared to WT hESCs (See Table 2 and Figure 4):

- ^13^C_4_-Glu/^13^C_3_-Lac (p=0.05),
- ^13^C_2_-Glu/^13^C_3_-Lac (p=0.009),
- ^13^C_3_-Glu/^13^C_3_-Lac (p=0.008)
- ^13^C_3_/^13^C_2_{^13^C_3_}-Lac (p=0.10).

**Table 2:** Metabolic flux ratios and 13C Enrichment for WT and LIS1 hESCs cultured in standard media (n=4). Additionally, matched ratios between LIS1 and WT hESCs cultured in parallel are shown to highlight relative changes within each batch.

|  |  |  |  |  |  |  |  |  |  |
| --- | --- | --- | --- | --- | --- | --- | --- | --- | --- |
|  |  | Total Metabolic Profile |  |  |  |  | Matched Ratios |  |  |
|  |  | WT |  | LIS |  | p value * | LIS/WT |  | p value * |
| Intermolecular<br>Metabolic Flux Ratio | $^{13}\text{C}_1\text{-Glc}/^{13}\text{C}_3\text{-Lac}$ | 1.46 ± | 0.14 | 1.44 ± | 0.19 | | 1.08 ± | 0.20 | |
| | $^{13}\text{C}_2\text{-Ac}/^{13}\text{C}_3\text{-Lac}$ | 0.053 ± | 0.033 | 0.072 ± | 0.048 | | 1.29 ± | 0.17 | p=0.01 |
| | $^{13}\text{C}_3\text{-Ala}/^{13}\text{C}_3\text{-Lac}$ | 0.035 ± | 0.010 | 0.035 ± | 0.008 | | 1.04 ± | 0.27 | |
| | $^{13}\text{C}_3\text{-Ser}/^{13}\text{C}_3\text{-Lac}$ | 0.020 ± | 0.010 | 0.030 ± | 0.011 | | 1.80 ± | 1.08 | |
| | $^{13}\text{C}_3\text{-Pyr}/^{13}\text{C}_3\text{-Lac}$ | 0.019 ± | 0.015 | 0.019 ± | 0.021 | | 1.64 ± | 1.53 | |
| | $^{13}\text{C}_3\text{-Form}/^{13}\text{C}_3\text{-Lac}$ | 0.0036 ± | 0.0012 | 0.0033 ± | 0.0012 | | 0.99 ± | 0.44 | |
| | $^{13}\text{C}_4\text{-Glu}/^{13}\text{C}_3\text{-Lac}$ | 0.0074 ± | 0.0021 | 0.0121 ± | 0.0031 | p=0.05 | 1.73 ± | 0.68 | p=0.07 |
| | $^{13}\text{C}_2\text{-Glu}/^{13}\text{C}_3\text{-Lac}$ | 0.0020 ± | 0.0004 | 0.0037 ± | 0.0008 | p=0.009 | 1.91 ± | 0.52 | p=0.01 |
| | $^{13}\text{C}_3\text{-Glu}/^{13}\text{C}_3\text{-Lac}$ | 0.0015 ± | 0.0003 | 0.0024 ± | 0.0004 | p=0.008 | 1.74 ± | 0.52 | p=0.03 |
| Intramolecular<br>Metabolic Flux<br>Ratios | $^{13}\text{C}_1/^{13}\text{C}_3\text{ Lac}$ | 61 ± | 4 | 65 ± | 5 | | 0.97 ± | 0.05 | |
| | $^{13}\text{C}_2\{^{12}\text{C}_3\}/^{13}\text{C}_3\text{ Lac}$ | 61 ± | 4 | 66 ± | 4 | | 0.95 ± | 0.07 | |
| | $^{13}\text{C}_2\{^{13}\text{C}_3\}/^{13}\text{C}_3\text{ Lac}$ | 95 ± | 6 | 116 ± | 20 | p=0.10 | 0.90 ± | 0.08 | p=0.04 |
| | $^{13}\text{C}_4/^{13}\text{C}_2\text{ Glu}$ | 3.8 ± | 0.6 | 3.3 ± | 0.1 | | 0.88 ± | 0.13 | |
| | $^{13}\text{C}_4/^{13}\text{C}_3\text{ Glu}$ | 5.1 ± | 0.9 | 4.9 ± | 0.7 | | 0.97 ± | 0.16 | |
| | $^{13}\text{C}_2/^{13}\text{C}_3\text{ Glu}$ | 1.4 ± | 0.2 | 1.5 ± | 0.1 | | 1.10 ± | 0.17 | |
| Relative<br>$^{13}\text{C}$ Enrichment | $^{13}\text{C}_3/^{13}\text{C}_2\text{-Ala}$ | 28 ± | 3 | 22 ± | 4 | | 0.80 ± | 0.21 | p=0.09 |
| | $^{13}\text{C}_3/^{13}\text{C}_2\text{-Ser}$ | 11 ± | 1 | 12 ± | 5 | | 1.06 ± | 0.44 | |
| Absolute<br>$^{13}\text{C}$ Enrichment | Frac $^{13}\text{C}_1\text{-Glc}$ | 0.98 ± | 0.01 | 0.98 ± | 0.01 | | 1.00 ± | 0.01 | |
| | Frac $^{13}\text{C}_3\text{-Lac}$ | 0.42 ± | 0.01 | 0.42 ± | 0.01 | | 0.99 ± | 0.02 | |
| | Frac $^{13}\text{C}_2\text{-Ac}$ | 0.27 ± | 0.10 | 0.25 ± | 0.12 | | 0.90 ± | 0.15 | |
| | Frac $^{13}\text{C}_1\text{-Form}$ | 0.12 ± | 0.02 | 0.11 ± | 0.02 | | 0.90 ± | 0.07 | p=0.03 |
| *For simplicity only p<0.1 are shown |  |  |  |  |  |  |  |  |  |

**Figure 4:**
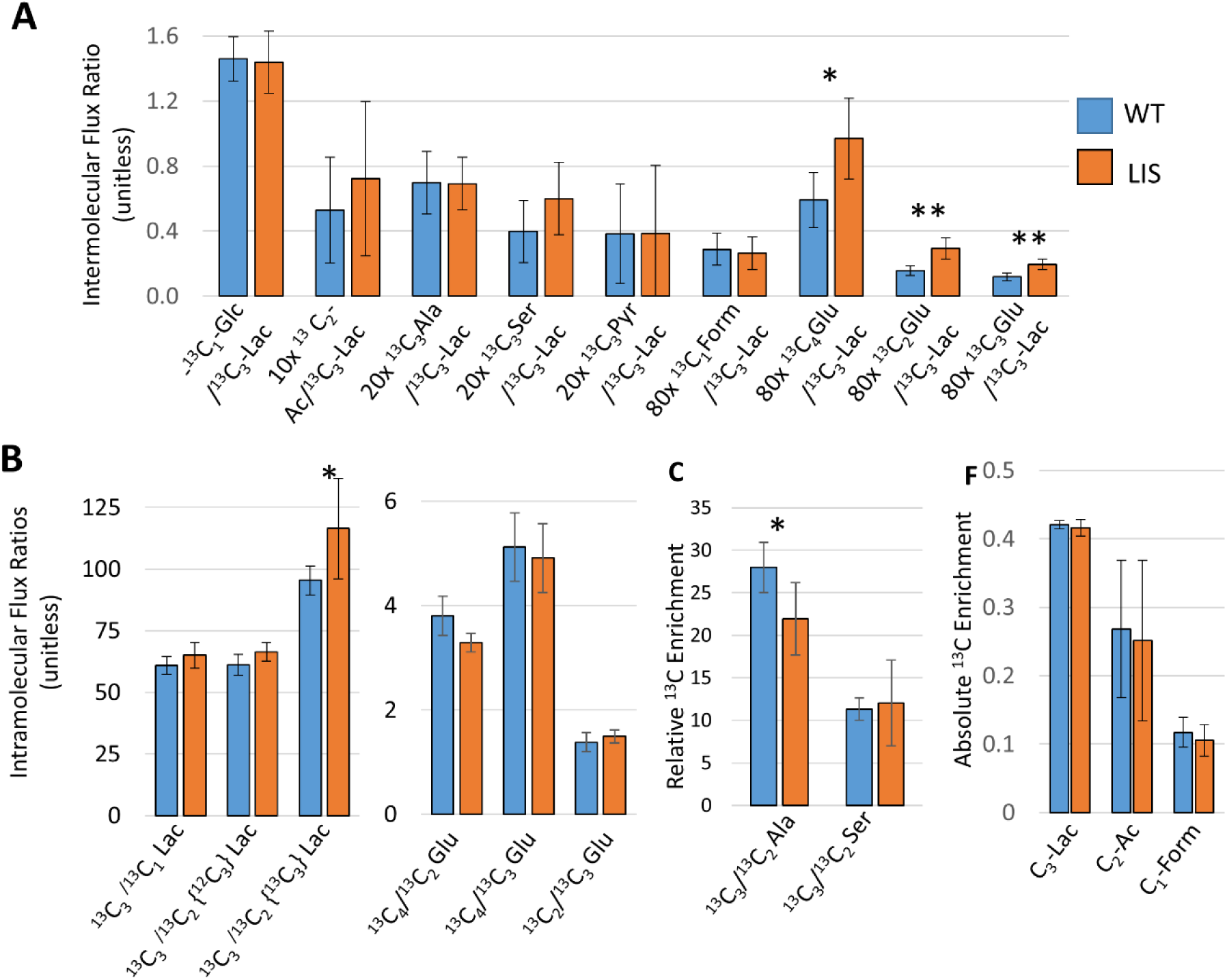
Metabolic fingerprint of WT (n=4) and LIS1 mutant (n=4) hESCs cultured in standard media. (A) Intermolecular flux ratios, (B) intramolecular flux ratios (C) relative ^13^C enhancements sand (D) absolute ^13^C enrichments. * p<0.1; ** p<0.01, ***p<0.005

These elevations suggest an enhanced routing of glucose-derived carbon through the TCA cycle and increased production of glutamate in the LIS1-deficient background.

Similar trends were observed for LIS1 mutant hESCs in media with L-Gln omission; however, these didn’t reach statistical significance (See SI Table S2).

To assess whether more subtle, genotype-specific differences in metabolic flux are masked by inter-batch variability, we performed matched pair analyses – directly comparing flux ratios and ^13^C enrichment values between LIS1 mutant and WT hESCs cultured in parallel. This approach recapitulated the previously observed increases in intermolecular flux ratios (See Table 2), specifically in:

- ^13^C_4_-Glu/^13^C_3_-Lac (p=0.07),
- ^13^C_2_-Glu/^3^C_3_-Lac (p=0.01),
- ^13^C_3_-Glu/^13^C_3_-Lac (p=0.03)
- ^13^C_3_/ ^13^C_2_{^13^C_3_}-Lac (p=0.04).

Importantly, matched analysis revealed additional significant metabolic differences not previously detected, specifically an increase in the ^13^C_2_-Ac/^13^C_3_-Lac metabolic flux ratio for LIS hESCs (p = 0.01). These findings suggest that *LIS1* haploinsufficiency affects not just glutamate biosynthesis but also other pathways connected to the TCA cycle, namely acetate production. In addition, from matched ratios, the following significant changes in ^13^C enrichment were observed:

- A decrease in the ^13^C_3_/^13^C_2_-Ala ratio (p=0.09)
- A decrease in the absolute ^13^C enrichment of C_1_-Form (p=0.03).

Similar trends were observed in media with L-Gln omission; further supporting the robustness of the metabolic phenotype (Supplementary Table S3).

### Interpreting Absolute and Relative ^13^C Enrichment

The observed ^13^C enrichment (*f_obs_*), whether absolute as measured from ^1^H NMR or relative as measured from ^13^C NMR, reflects both the fractional ^13^C label of newly formed product (*f_met_*) and dilution by pre-existing unlabeled pools. For lactate, no detectable ^1^H signal is observed in media incubated without hESCs, indicating negligible background. Therefore, the fractional ^13^C labeling of C_3_-Lactate can be assumed as directly equal to the observed labeling: *f_met_=f_obs_*. In contrast, for metabolites like acetate and formate, which are present in the culture media even in the absence of cells, isotopic dilution must be accounted for. The fractional metabolic labeling (*f_met_*) can be derived by fitting the following equation that incorporated natural abundance (1%) background ^13^C labeling:

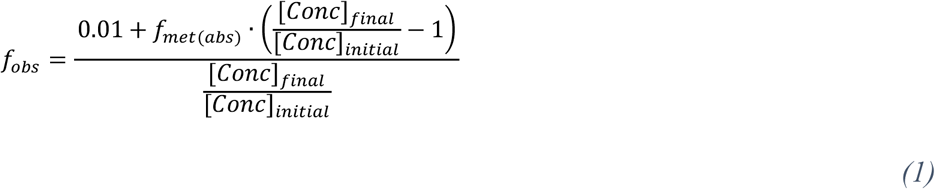

Using this approach, we calculated:

- For C_2_-Acetate: *f_met(abs)_* = 0.43±0.01 (R^2^=0.94) (Figure 5A)
- For C_1_-Formate *f_met(abs)_* = 0.17±0.02 (R^2^=0.58) (Figure 5B).

**Figure 5:**
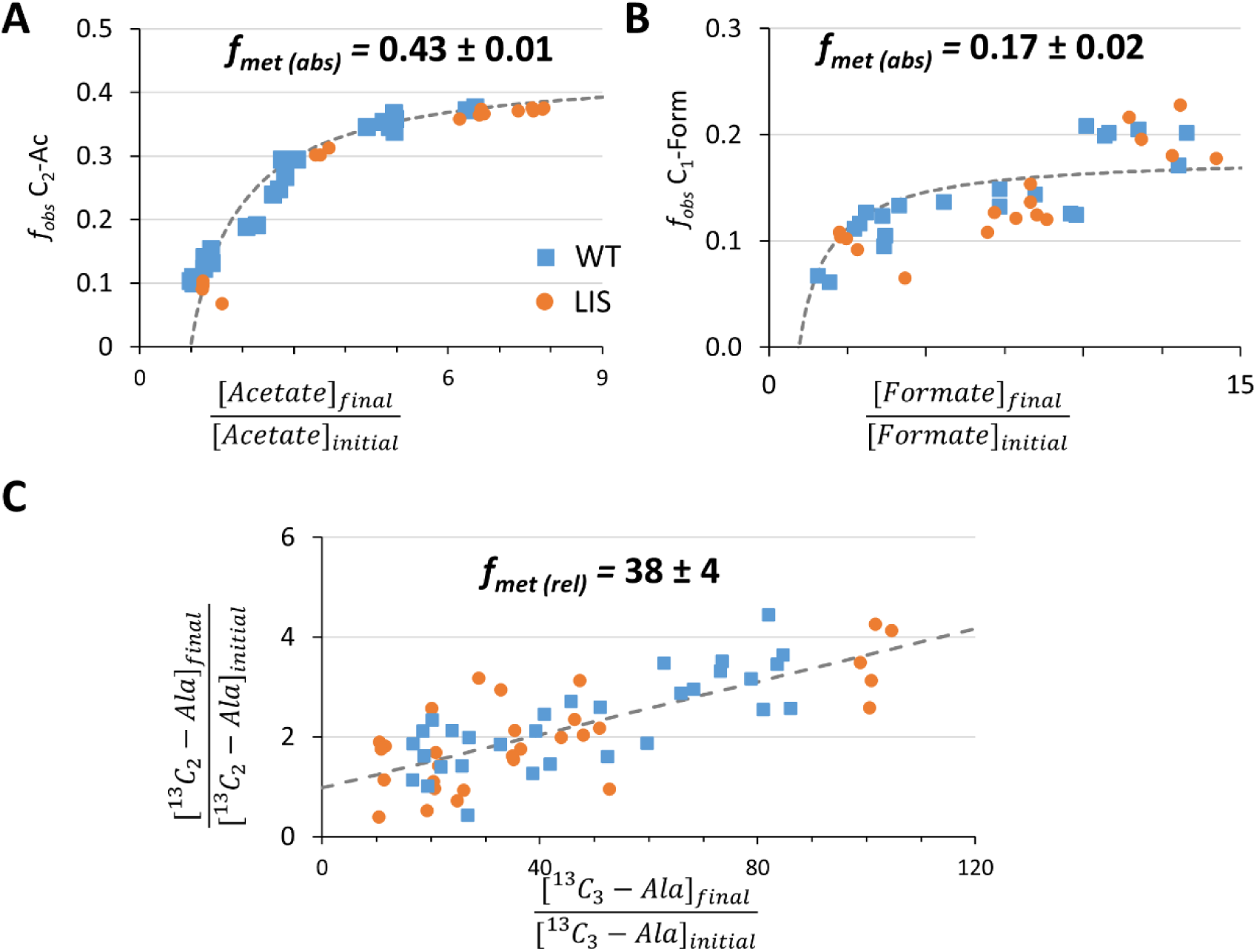
Determining the fractional ^13^C enrichment of metabolic products (*f_met_*). (A-B) Absolute ^13^C enrichment (*f_met(abs)_*) of C_2_-Acetate and C_1_-Formate, determined by fitting the observed ^13^C enrichment as function of the change in metabolite concentration using Eq. 1. (C) Relative ^13^C enrichment (*f_met(rel)_*) of C_3_-Alanine, determined by fitting the fold change in C_2_ signal vs. the fold change in C_3_ signal using Eq. 2. Together, these measurements quantify the proportion of each metabolite pool derived from ^13^C-labeled glucose.

We also examined relative metabolic labeling for alanine. By comparing the fold-change in C_2_ versus C_3_ signal intensities, we can gain insight into the relative fractional metabolic labeling of C_3_/C_2_-Alanine by fitting it to the following equation:

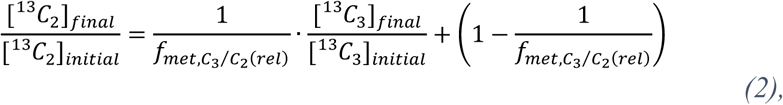

where 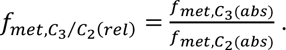

This yielded *f_met,C3/C2-Ala(rel)_*=38 ± 4 (R^2^=0.58). Assuming negligible ^13^C enrichment of the C_2_-Ala position, this corresponds to *f_met,C3-Ala(abs)_*= 0.42 ± 0.04, remarkably consistent with that of C_2_-acetate and C_3_-lactate.

## Discussion

### Metabolic phenotype of hESCs

Using the Intelliwaste method, we confirmed the characteristic glycolytic metabolic phenotype previously reported in naïve hESCs ^33^ using mass spectrometry. Specifically, among the diverse substrates present in the nutrient-rich culture media, glucose is the only one significantly consumed, and the majority of this glucose is converted to Lactate. In our data, 68 ± 7 % of the consumed ^13^C_1_-Glucose label is recovered as ^13^C_3_-Lactate.

The observed fractional ^13^C enrichment of the C_3_-Lactate site was 0.42 ± 0.01, compared to a maximum theoretical enrichment of 0.49 (accounting for the 0.98 ± 0.01 fractional enrichment of ^13^C_1_-Glucose). From this, we infer that 86% of this Lactate is indeed produced from glucose via glycolysis. The remaining 14% may reflect the contribution of other substrates to the glycolytic intermediates upstream to Lactate, or metabolism of Glucose via the Pentose Phosphate Pathway (PPP), where the ^13^C_1_ label is lost as ^13^CO_2_. Future experiments using ^13^C_6_-Glucose labeling will be instrumental in distinguishing between PPP activity and alternate substrate contributions to lactate production.

Metabolic flux observed at a ^13^C metabolite site does not necessarily imply direct metabolic conversion from the ^13^C_1_-Glucose precursor, as ^13^C has 1.1% natural abundance. The fractional ^13^C enrichment of 0.42 ± 0.01 observed for ^13^C_3_-Lac reflects specific isotopic enrichment from the labeled glucose. In contrast, other lactate sites didn’t show significant enrichment beyond natural abundance. Based on *a priori* knowledge of biochemical pathways, we interpret the flux observed at the ^13^C_3_-Glutamate site primarily as an increase in metabolite concentration combined with the 1.1% natural ^13^C abundance, rather than incorporation of ^13^C label from ^13^C_1_-Glucose.

Seven additional metabolite sites exhibited clear specific isotopic enrichment, including ^13^C_2_-Ac, ^13^C_3_-Ala, ^13^C_3_-Ser, ^13^C_3_-Pyr, ^13^C_1_-Form, ^13^C_2_-Glu and ^13^C_4_-Glu. Together, these accounted for approximately 10 ± 5% of the consumed ^13^C_1_-Glucose label. The remaining 22 ± 12% of the ^13^C_1_-Glucose signal was either converted to ^13^CO_2_ (which exchanges with ambient air and is lost during the incubation process), incorporated into biomass and intracellular metabolite pools, or was below the detection threshold of our NMR measurements.

The fractional metabolic labeling of ^13^C_2_-Ac (0.43 ± 0.01) and ^13^C_3_-Alanine (0.45 ± 0.09, assuming natural abundance enrichment of C_2_-Alanine) closely match that of ^13^C_3_-Lactate. This result indicates that in hESCs, both ^13^C_2_-Acetate and ^13^C_3_-Alanine are predominantly produced from the same intracellular pyruvate pool as ^13^C_3_-Lactate. In contrast, the fractional metabolic labeling of ^13^C_1_-Form is significantly lower (0.17 ± 0.01), indicating that in addition to serine synthesized from the glycolytic intermediate 3-Phosphoglycerate, other unlabeled substrates substantially contribute to the intracellular serine pool (See Figure 6A). The close similarity between the observed isotopic enrichment of newly produced ^13^C_1_-Formate (0.13 ± 0.04) and that of C_3_-Serine (0.12 ± 0.02, assuming natural abundance C_2_-Ser enrichment) further suggests that an equilibrium is rapidly established between the intracellular and extracellular C_3_-Ser pools. Consequently, the isotopic enrichment of this combined pool likely determines the fractional metabolic labeling of ^13^C_1_-Formate. This result may explain why the ^13^C_3_-Ser flux does not scale proportionally with the ^13^C_3_-La flux across samples, but instead remains relatively constant in absolute concentrations across different batches (See Figure 2B-C and Supplementary Figures S1-S2).

**Figure 6:**
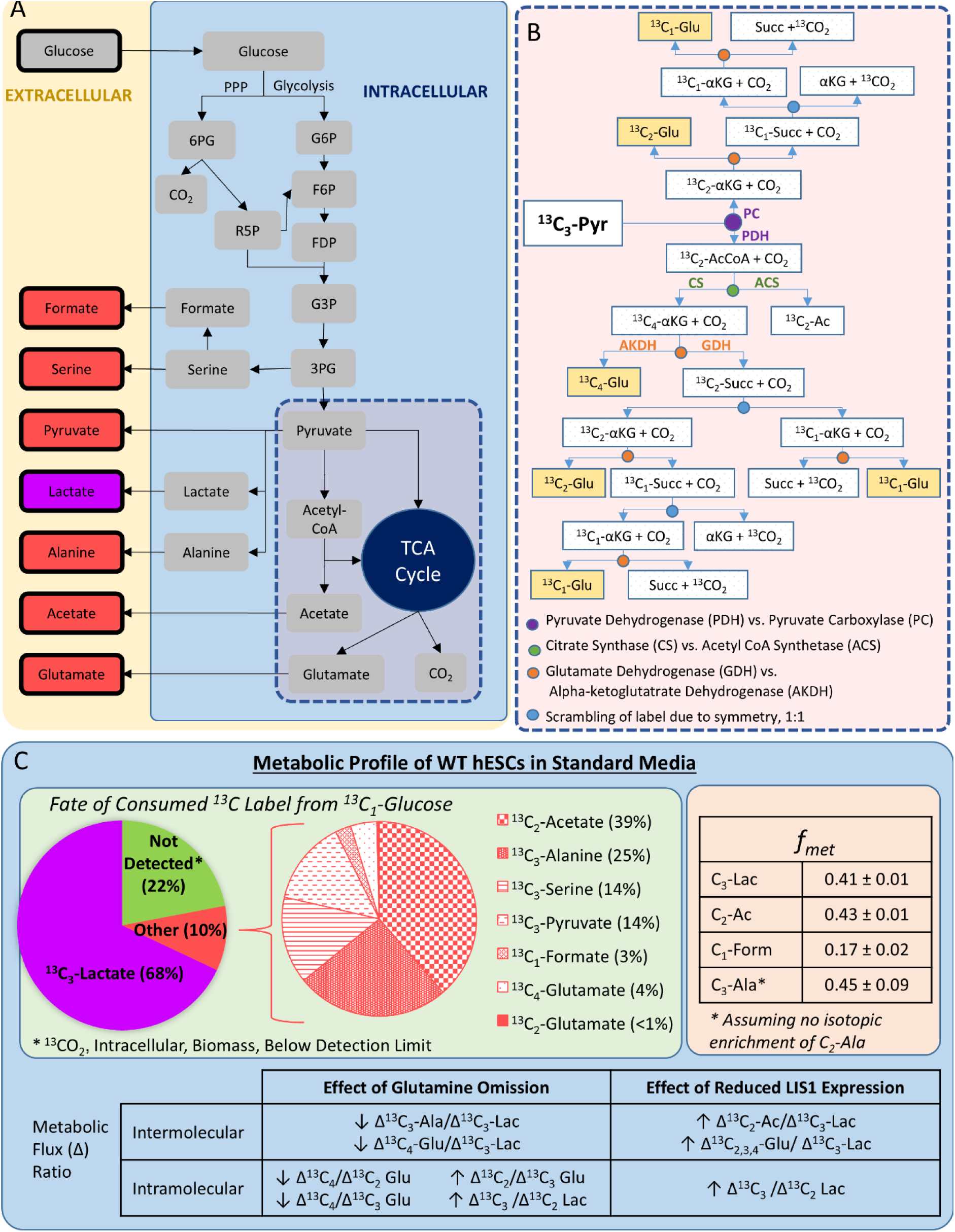
Graphical summary of insights into glucose metabolism of hESCs from Intelliwaste measurements. (A) Schematic of ^13^C_1_-Glucose metabolism by hESCs. Metabolic products detected in extracellular media after 24 hours of incubation are color-coded: Lactate (purple) and additional ^13^C enriched products (red). (B) Schematic depiction of how the different enzymes of the TCA cycle affect the isotopic enrichment of the different sites of Glutamate. (C) Integrated metabolic fingerprint of hESCs derived from Intelliwaste analysis. Top panels summarize flux and enrichment data for WT hESCs cultured in standard media; the lower table highlights significant changes observed in response to glutamine omission and when comparing LIS1 mutant to WT hESCs.

Glutamate is produced via the conversion of the TCA cycle intermediate α-Ketoglutarate to Glutamate. The relative isotopic enrichment of Glutamate provides insight into the activity of intracellular enzymes, as ^13^C_3_-Pyr can enter the TCA cycle by two distinct routes: via Pyruvate Carboxylase (PC), which leads to enrichment of ^13^C_2_-Glutamate, or via Pyruvate Dehydrogenase (PDH), leading to preferential labeling of ^13^C_4_-Glu (see Figure 6B). In hESCs cultured with Glutamine, the observed ^13^C_4_/^13^C_2_ Glutamate flux ratio of 3.8 ± 0.6 indicated that PDH activity substantially exceeds that of PC. In contrast, under glutamine-depleted conditions, this ratio is reduced by approximately two-fold, suggesting a relative decrease in the PDH/PC activity ratio, although PDH activity is still higher than PC activity. This conclusion is further supported by the opposing trends in isotopic enrichment: ^13^C_2_ Glutamate enrichment increases, while ^13^C_4_ Glutamate enrichment decreases in the absence of glutamine.

Moreover, omitting L-Glutamine reduces the activity of Alanine aminotransferase (ALT) enzyme, as evidenced by the approximately two-fold reduction in the ^13^C_3_-Alanine/^13^C_3_-Lactate flux ratio. Interestingly, assuming ^13^C_3_-Glu is not specifically enriched, the overall glutamate-to-lactate metabolic flux ratio remains largely unchanged upon glutamine withdrawal. This suggests that while glutamine availability influences specific TCA cycle enzymatic pathways, it does not dramatically affect the net flux to glutamate.

Notably, these mechanistic insights into enzyme activity and carbon routing would not be accessible without the isotopic labeling enabled by the Intelliwaste method. For example, the highest observed isotopic enrichment in Glutamate site - belonging to C_4_-Glutamate in the presence of Glutamine - is only 5.1 ± 0.9 fold times above natural abundance (assuming C_3_-Glu is at natural abundance enrichment). This level of enrichment is considerably lower than what is observed for C_3_-Lac, C_3_-Ala and C_2_-Ac. The modest enrichment in glutamate cannot be attributed solely to dilution by unlabeled glutamine-derived glutamate, since similar trends are observed even when Glutamine is omitted from the media. Nor is it likely due to preferential routing of pyruvate carbons toward oxidative phosphorylation rather than glutamate biosynthesis, as such a shift would affect glutamate production relative to CO₂ but not its isotopic enrichment. Instead, this is a clear indication that in addition to Pyruvate derived from Glucose other substrates are metabolized via the TCA cycle by hESCs.

The metabolic fingerprint of the *LIS1* mutant (LIS) hESCs, measured under both L-Glutamine-replete and depleted conditions, was largely similar to that of wild-type (WT) hESCs – consistent with the expectation that reduced expression of LIS1, a protein not directly involved in metabolic pathways, would not drastically alter global metabolism. Nevertheless, several statistically significant differences were observed. Notably, LIS hESCs showed increased C_2,3,4_-Glu/C_3_-Lac metabolic flux ratios, suggesting elevated glutamate production. When comparing matched samples, LIS hESCs also exhibited a significantly higher ^13^C_2_-Ac/^13^C_3_-Lac ratio, indicating enhanced acetate output. In addition, the ^13^C_4_/^13^C_2_-Glu metabolic flux ratio was significantly decreased in LIS hESCs, indicating a decrease in the PDH/PC activity ratio and thus a relative shift in TCA cycle entry route preference compared to WT hESCs.

To further dissect these metabolic shifts, we plan to enrich the growth media with ^13^C-labeled amino acids and/or omit specific amino acids in future studies. This will help determine the contributions of individual substrates to the TCA cycle more precisely in both WT and LIS hESCs. In addition, using media enriched with doubly labeled ^13^C_1,2_-Glucose will enable us to evaluate the relative activity of the TCA cycle enzymes, such as Glutamate Dehydrogenase and Alpha-Ketoglutarate Dehydrogenase (See Figure 6, right panel).

Beyond defining general metabolic fingerprints, the Intelliwaste method also enables the identification of batches with aberrant metabolic behavior. For example, Batch 5 exhibited markedly reduced metabolic flux ratios for ^13^C_2_-Ac/^13^C_3_-Lac and ^13^C_3_-Pyr/^13^C_3_-Lac, across both WT and LIS hESCs (See Figure 2C-D and Supplementary Figure S2). Previous studies have linked decreased acetate metabolism to early differentiation of hESCs ^34^, suggesting that Intelliwaste may serve as a sensitive tool for detecting early, inadvertent loss of pluripotency in naïve hESC cultures - a common challenge in stem cell maintenance.

### Expanding the Intelliwaste Method

The Intelliwaste method offers a powerful and non-destructive approach for characterizing complex metabolic phenotypes by isotopically labeling extracellular metabolites. As demonstrated here for naïve hESCs, Intelliwaste can capture subtle metabolic differences between genetically normal and mutated cells. One of the key advantages of this method lies in its focus on extracellular fluxes, which yield metabolite concentrations several orders of magnitude higher than those found intracellularly. For example, based on the results of Fan et. al. ^35^ for an immortal mammalian cell line, the amount of lactate secreted over a 24-hour incubation is approximately 100-fold greater than the concentration of the most abundant intracellular glycolytic or TCA cycle intermediates.

The non-invasive nature of Intelliwaste enables repeated sampling from the same culture, making it suitable for time-course experiments as well as for parallel integration with orthogonal assays. In this study, we employed a relatively simple implementation of Intelliwaste, using only a single labeled substrate (^13^C_1_-glucose) and altering a single experimental variable (the presence or absence of L-glutamine). Even under these minimal perturbations, the method was able to generate a detailed metabolic fingerprint and detect statistically significant effects of the LIS1 mutation. The strong reproducibility across replicate plates suggests that Intelliwaste is highly amenable to multiplexed experimental designs, where media composition and/or isotopic enrichment are systematically varied.

Looking ahead, the method’s capacity for longitudinal monitoring of stem cell metabolism, from naïve hESCs through differentiation into mature brain organoids, opens the door to tracing how early-stage metabolic phenotypes, such as those associated with *LIS1* deficiency, influence long-term outcomes in organoid development and function.

## Materials and Methods

### Cell Culture and Ethical Statement

An NIH-approved hESC line, NIHhESC-10-0079, WIBR3 (W3), was used in this study. Isogenic mutant cell-line clones were previously generated by CRISPR-Cas9-mediated heterozygous deletion in the LIS1 gene ^19^. Work with hESC (WIBR3, NIHhESC-10-0079) and genome editing was carried out with approval from the Weizmann Institute of Science IRB (Institutional Review Board). All cell lines were regularly checked for mycoplasma contamination. hESCs were cultured in naïve media ^32^ on 1% Matrigel-coated 6-well plates, seeded at 70,000 cells per well, and maintained in 2.5 ml of media, which was refreshed every 24 hours. For 1-^13^C labeled experiments, naïve media was prepared as above, except the standard DMEM-F12 was replaced with SILAC Advanced DMEM/F-12 Flex Media (ThermoFisher). To match standard DMEM-F12 composition, L-arginine, L-lysine, and phenol red were added. Additionally, 1.81 g/L of 1-^13^C Glucose (Cambridge Isotopes) was included, yielding a final glucose concentration of 10 mM with >99% isotopic enrichment at the ^13^C_1_ position.

For glutamine omission experiments, the 1-^13^C Glucose media was prepared without the standard L-Glutamine supplement.

In Intelliwaste experiments, cells were rinsed twice with 3 mL of PBS and incubated for 24 hours in 2 mL of 10 mM 1-^13^C Glucose naïve media, with or without L-Glutamine. After 24 hours, the media was collected and stored in Eppendorf tubes at -80 ^°^C. As a control, media was incubated in plates without cells under identical conditions.

### NMR Measurements

Aliquots of spent media, incubated with and without hESCs, were defrosted and mixed in a 6.25:1 ratio with a standard solution of D_2_O containing sodium azide and dioxane (used as a chemical shift reference). NMR spectra were acquired on a Bruker AVANCE NEO-600 NMR spectrometer equipped with a 5 mm TCI-xyz CryoProbe at 298 K. ^1^H spectra were acquired with water suppression and under two conditions:

- With ^13^C decoupling (decoupling on ^13^C channel centered at 50 ppm)
- Without ^13^C decoupling (decoupling on ^13^C channel centered at 400 ppm).

Acquisition parameters included spectral width (sw) of 20 ppm and a delay (d1) of 4 seconds. ^13^C NMR spectra were acquired using a DEPTQ sequence with ^1^H decoupling during acquisition, sw of 200 ppm and repetition time (TR) of 4 seconds.

In addition, for selected samples, standard pulse acquire ^13^C NMR spectra were acquired with ^1^H decoupling, sw of 237 ppm and an extended recycle delay (TR>20 seconds).

### Processing NMR spectra

NMR spectra were processed and analyzed using TOPSPIN 4.0 (Bruker BioSpin, Germany). For ^1^H spectra, data were zero-filled to 128k points, processed with a 0.10 Hz line broadening and zero- and first-order phase correction. For ^13^C spectra, the same zero filling was applied, with a 0.50 Hz line broadening and zero- and first-order phase correction.

### Metabolite quantification from ^13^C NMR spectra

Inspection of the ^13^C spectra showed noticeable variability in metabolite chemical shifts, attributable to small differences in sample pH. Although pH fluctuations were minor, they were sufficient to cause measurable shifts. To correct for this, we implemented a linear correction procedure based on the chemical shift of bicarbonate, a pH-sensitive but spectrally isolated peak (See Supplementary Table S1 and Supplementary Figure S1). For each metabolite site, the chemical shift was linearly adjusted relative to the shift of bicarbonate.

Peak fitting was performed using custom-written scripts in Matlab (Mathworks), applying Lorentzian line shape models to 100 unique ^13^C NMR peaks. To improve the accuracy of the fitting process, peaks with chemical shift separation <4 Hz were excluded. This approach enabled successful fitting of 96% of metabolite peaks across the 97 acquired ^13^C NMR spectra. To reduce redundancy, C_6_ glucose peaks were modeled based on the position of the C_1_ glucose peak, reducing the number of independent peaks to 98 and increasing the fitted success rate to 98%. Since dioxane was included as an internal reference in all samples, the final number of independent metabolite sites was 97.

Metabolite fluxes were determined by subtracting the median peak integral in the spent media (from hESCs incubated wells) from that in control media (incubated without hESCs). To convert these fluxes from signal intensity units to concentrations, a correction factor, *c*, was calculated for each metabolite peak. This factor was derived as the ratio of the signal integral acquired using a fully relaxed ^13^C-quantitative NMR spectrum (acquired without NOE, denoted as *Signal Intensity*_13*C*−*quant*_) to that from a ^13^C-DEPT spectrum (*Signal Intensity*_13*C*−*DEPT*_).

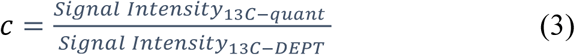

### Metabolite Concentration and ^13^C Enrichment from ^1^H NMR

For formate and acetate, we determined the fold-change in metabolite concentration by comparing the DMSO normalized peak intensity of the peak in spent media compared to fresh media, with the fold concentration increase given as:

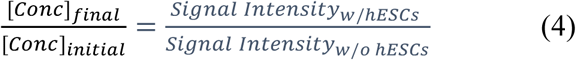

For formate, acetate, lactate and glucose we determined the observed fraction of ^13^C label (*f_obs_*) in the C_1_, C_2_, C_3_ and β-C_1_ positions, respectively, by comparing the intensity of the central ^1^H{^12^C} peak to the satellite ^1^H{^13^C} peaks.

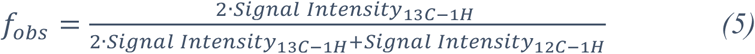

### Statistical Analysis

Statistical analysis was performed using a two-tailed Student’s t-test assuming equal variance, implemented in Microsoft Excel. Where applicable, curve fitting, as well as calculation of confidence intervals and R² values, was performed using MATLAB (MathWorks).

## Data availability

Spectrum data is available at https://doi.org/10.34933/aa7b42fa-d76a-457a-8fd8-b39a4aa3b7b2.

## Supplementary material

Supplementary material is included in a separate PDF file.

---

**Table S1:**
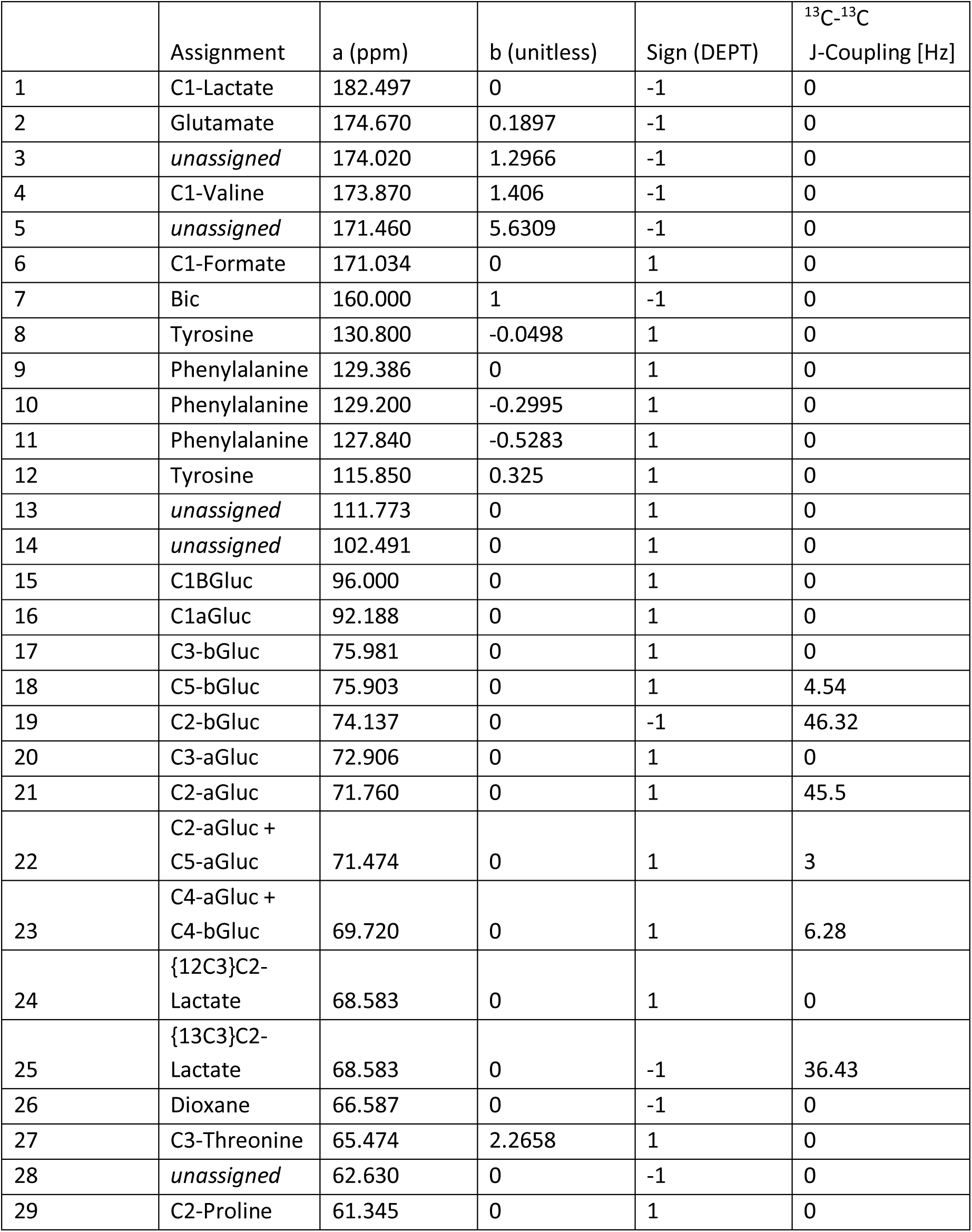
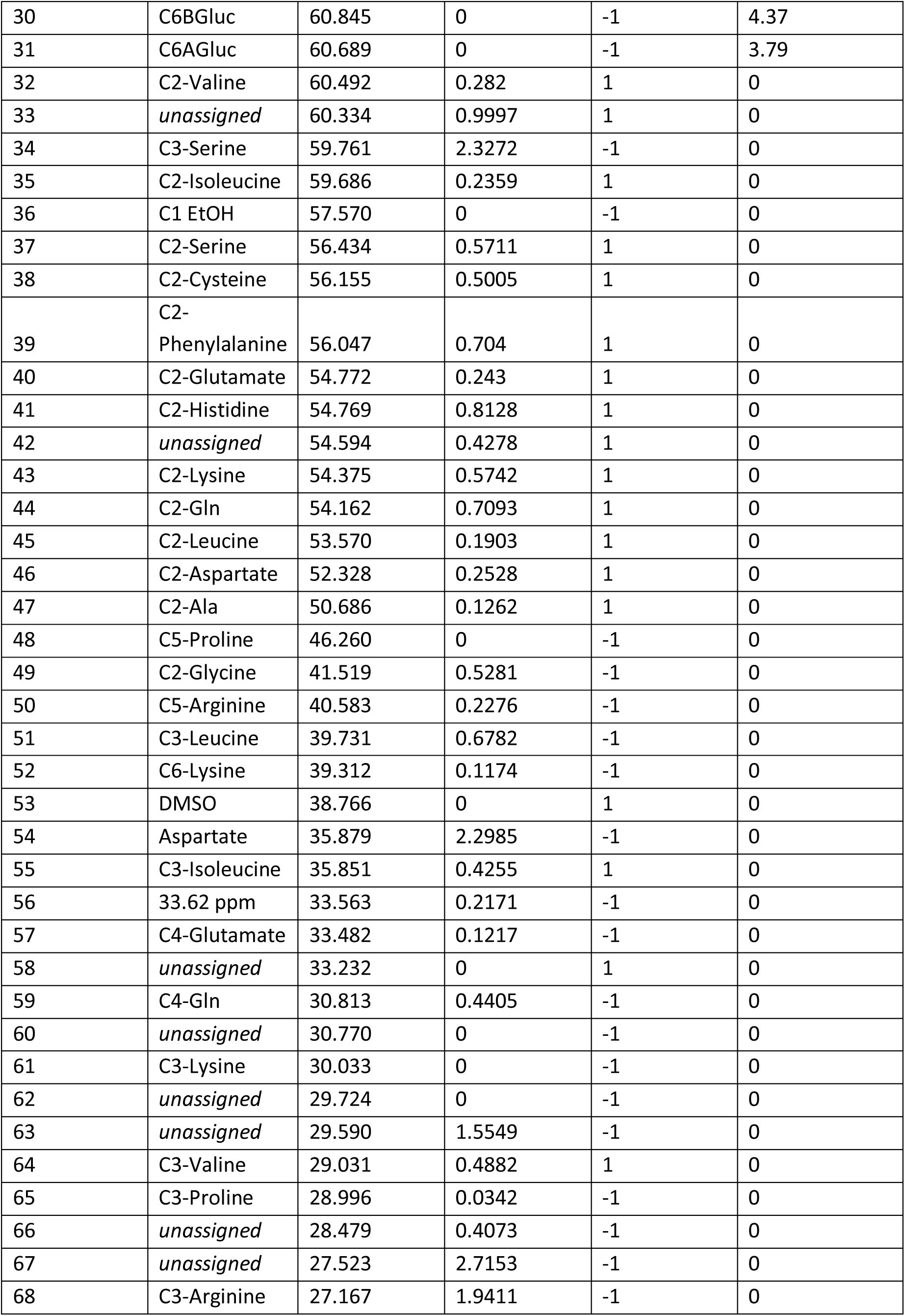
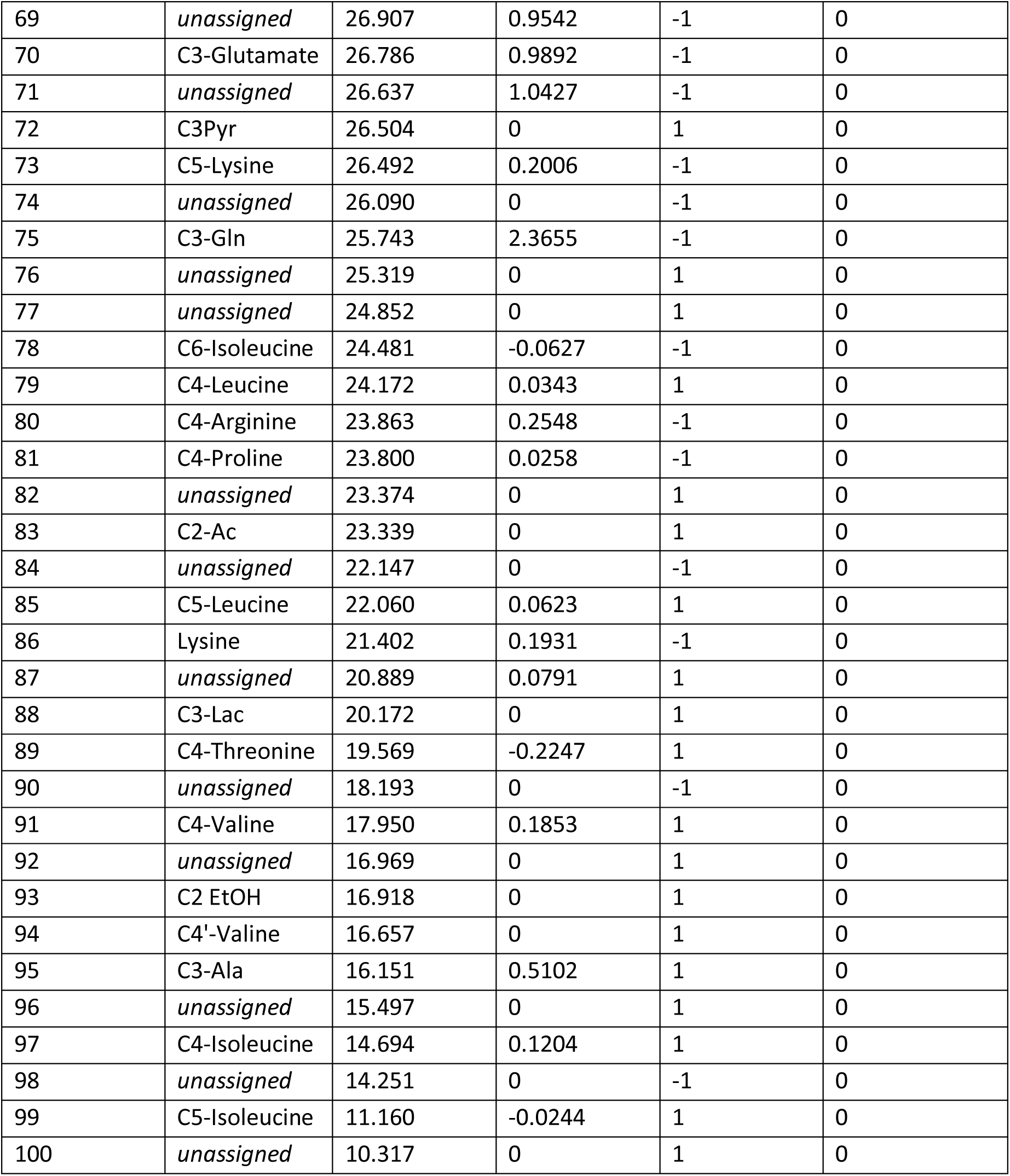
List of peaks used to fit the ^13^C NMR data. A linear equation was used to model the pH-dependent chemical shift of each metabolite over the observed range of bicarbonate chemical shifts (160.2-160.6 ppm) : δ_*met*_ = *a* + *b* · (δ_*Bic*_ − 160 *ppm*). Values of the slope (*b*) and intercept (*a*) for each peak are provided in the table.

| | Assignment | a (ppm) | b (unitless) | Sign (DEPT) | $^{13}\text{C}$-$^{13}\text{C}$<br>J-Coupling [Hz] |
| --- | --- | --- | --- | --- | --- |
| 1 | C1-Lactate | 182.497 | 0 | -1 | 0 |
| 2 | Glutamate | 174.670 | 0.1897 | -1 | 0 |
| 3 | *unassigned* | 174.020 | 1.2966 | -1 | 0 |
| 4 | C1-Valine | 173.870 | 1.406 | -1 | 0 |
| 5 | *unassigned* | 171.460 | 5.6309 | -1 | 0 |
| 6 | C1-Formate | 171.034 | 0 | 1 | 0 |
| 7 | Bic | 160.000 | 1 | -1 | 0 |
| 8 | Tyrosine | 130.800 | -0.0498 | 1 | 0 |
| 9 | Phenylalanine | 129.386 | 0 | 1 | 0 |
| 10 | Phenylalanine | 129.200 | -0.2995 | 1 | 0 |
| 11 | Phenylalanine | 127.840 | -0.5283 | 1 | 0 |
| 12 | Tyrosine | 115.850 | 0.325 | 1 | 0 |
| 13 | *unassigned* | 111.773 | 0 | 1 | 0 |
| 14 | *unassigned* | 102.491 | 0 | 1 | 0 |
| 15 | C1BGluc | 96.000 | 0 | 1 | 0 |
| 16 | C1aGluc | 92.188 | 0 | 1 | 0 |
| 17 | C3-bGluc | 75.981 | 0 | 1 | 0 |
| 18 | C5-bGluc | 75.903 | 0 | 1 | 4.54 |
| 19 | C2-bGluc | 74.137 | 0 | -1 | 46.32 |
| 20 | C3-aGluc | 72.906 | 0 | 1 | 0 |
| 21 | C2-aGluc | 71.760 | 0 | 1 | 45.5 |
| 22 | C2-aGluc +<br>C5-aGluc | 71.474 | 0 | 1 | 3 |
| 23 | C4-aGluc +<br>C4-bGluc | 69.720 | 0 | 1 | 6.28 |
| 24 | {$^{12}\text{C}$3}C2-Lactate | 68.583 | 0 | 1 | 0 |
| 25 | {$^{13}\text{C}$3}C2-Lactate | 68.583 | 0 | -1 | 36.43 |
| 26 | Dioxane | 66.587 | 0 | -1 | 0 |
| 27 | C3-Threonine | 65.474 | 2.2658 | 1 | 0 |
| 28 | *unassigned* | 62.630 | 0 | -1 | 0 |
| 29 | C2-Proline | 61.345 | 0 | 1 | 0 |
| 30 | C6BGluc | 60.845 | 0 | -1 | 4.37 |
| 31 | C6AGluc | 60.689 | 0 | -1 | 3.79 |
| 32 | C2-Valine | 60.492 | 0.282 | 1 | 0 |
| 33 | *unassigned* | 60.334 | 0.9997 | 1 | 0 |
| 34 | C3-Serine | 59.761 | 2.3272 | -1 | 0 |
| 35 | C2-Isoleucine | 59.686 | 0.2359 | 1 | 0 |
| 36 | C1 EtOH | 57.570 | 0 | -1 | 0 |
| 37 | C2-Serine | 56.434 | 0.5711 | 1 | 0 |
| 38 | C2-Cysteine | 56.155 | 0.5005 | 1 | 0 |
| 39 | C2-Phenylalanine | 56.047 | 0.704 | 1 | 0 |
| 40 | C2-Glutamate | 54.772 | 0.243 | 1 | 0 |
| 41 | C2-Histidine | 54.769 | 0.8128 | 1 | 0 |
| 42 | *unassigned* | 54.594 | 0.4278 | 1 | 0 |
| 43 | C2-Lysine | 54.375 | 0.5742 | 1 | 0 |
| 44 | C2-Gln | 54.162 | 0.7093 | 1 | 0 |
| 45 | C2-Leucine | 53.570 | 0.1903 | 1 | 0 |
| 46 | C2-Aspartate | 52.328 | 0.2528 | 1 | 0 |
| 47 | C2-Ala | 50.686 | 0.1262 | 1 | 0 |
| 48 | C5-Proline | 46.260 | 0 | -1 | 0 |
| 49 | C2-Glycine | 41.519 | 0.5281 | -1 | 0 |
| 50 | C5-Arginine | 40.583 | 0.2276 | -1 | 0 |
| 51 | C3-Leucine | 39.731 | 0.6782 | -1 | 0 |
| 52 | C6-Lysine | 39.312 | 0.1174 | -1 | 0 |
| 53 | DMSO | 38.766 | 0 | 1 | 0 |
| 54 | Aspartate | 35.879 | 2.2985 | -1 | 0 |
| 55 | C3-Isoleucine | 35.851 | 0.4255 | 1 | 0 |
| 56 | 33.62 ppm | 33.563 | 0.2171 | -1 | 0 |
| 57 | C4-Glutamate | 33.482 | 0.1217 | -1 | 0 |
| 58 | *unassigned* | 33.232 | 0 | 1 | 0 |
| 59 | C4-Gln | 30.813 | 0.4405 | -1 | 0 |
| 60 | *unassigned* | 30.770 | 0 | -1 | 0 |
| 61 | C3-Lysine | 30.033 | 0 | -1 | 0 |
| 62 | *unassigned* | 29.724 | 0 | -1 | 0 |
| 63 | *unassigned* | 29.590 | 1.5549 | -1 | 0 |
| 64 | C3-Valine | 29.031 | 0.4882 | 1 | 0 |
| 65 | C3-Proline | 28.996 | 0.0342 | -1 | 0 |
| 66 | *unassigned* | 28.479 | 0.4073 | -1 | 0 |
| 67 | *unassigned* | 27.523 | 2.7153 | -1 | 0 |
| 68 | C3-Arginine | 27.167 | 1.9411 | -1 | 0 |
| 69 | *unassigned* | 26.907 | 0.9542 | -1 | 0 |
| 70 | C3-Glutamate | 26.786 | 0.9892 | -1 | 0 |
| 71 | *unassigned* | 26.637 | 1.0427 | -1 | 0 |
| 72 | C3Pyr | 26.504 | 0 | 1 | 0 |
| 73 | C5-Lysine | 26.492 | 0.2006 | -1 | 0 |
| 74 | *unassigned* | 26.090 | 0 | -1 | 0 |
| 75 | C3-Gln | 25.743 | 2.3655 | -1 | 0 |
| 76 | *unassigned* | 25.319 | 0 | 1 | 0 |
| 77 | *unassigned* | 24.852 | 0 | 1 | 0 |
| 78 | C6-Isoleucine | 24.481 | -0.0627 | -1 | 0 |
| 79 | C4-Leucine | 24.172 | 0.0343 | 1 | 0 |
| 80 | C4-Arginine | 23.863 | 0.2548 | -1 | 0 |
| 81 | C4-Proline | 23.800 | 0.0258 | -1 | 0 |
| 82 | *unassigned* | 23.374 | 0 | 1 | 0 |
| 83 | C2-Ac | 23.339 | 0 | 1 | 0 |
| 84 | *unassigned* | 22.147 | 0 | -1 | 0 |
| 85 | C5-Leucine | 22.060 | 0.0623 | 1 | 0 |
| 86 | Lysine | 21.402 | 0.1931 | -1 | 0 |
| 87 | *unassigned* | 20.889 | 0.0791 | 1 | 0 |
| 88 | C3-Lac | 20.172 | 0 | 1 | 0 |
| 89 | C4-Threonine | 19.569 | -0.2247 | 1 | 0 |
| 90 | *unassigned* | 18.193 | 0 | -1 | 0 |
| 91 | C4-Valine | 17.950 | 0.1853 | 1 | 0 |
| 92 | *unassigned* | 16.969 | 0 | 1 | 0 |
| 93 | C2 EtOH | 16.918 | 0 | 1 | 0 |
| 94 | C4'-Valine | 16.657 | 0 | 1 | 0 |
| 95 | C3-Ala | 16.151 | 0.5102 | 1 | 0 |
| 96 | *unassigned* | 15.497 | 0 | 1 | 0 |
| 97 | C4-Isoleucine | 14.694 | 0.1204 | 1 | 0 |
| 98 | *unassigned* | 14.251 | 0 | -1 | 0 |
| 99 | C5-Isoleucine | 11.160 | -0.0244 | 1 | 0 |
| 100 | *unassigned* | 10.317 | 0 | 1 | 0 |

**Figure S1:**
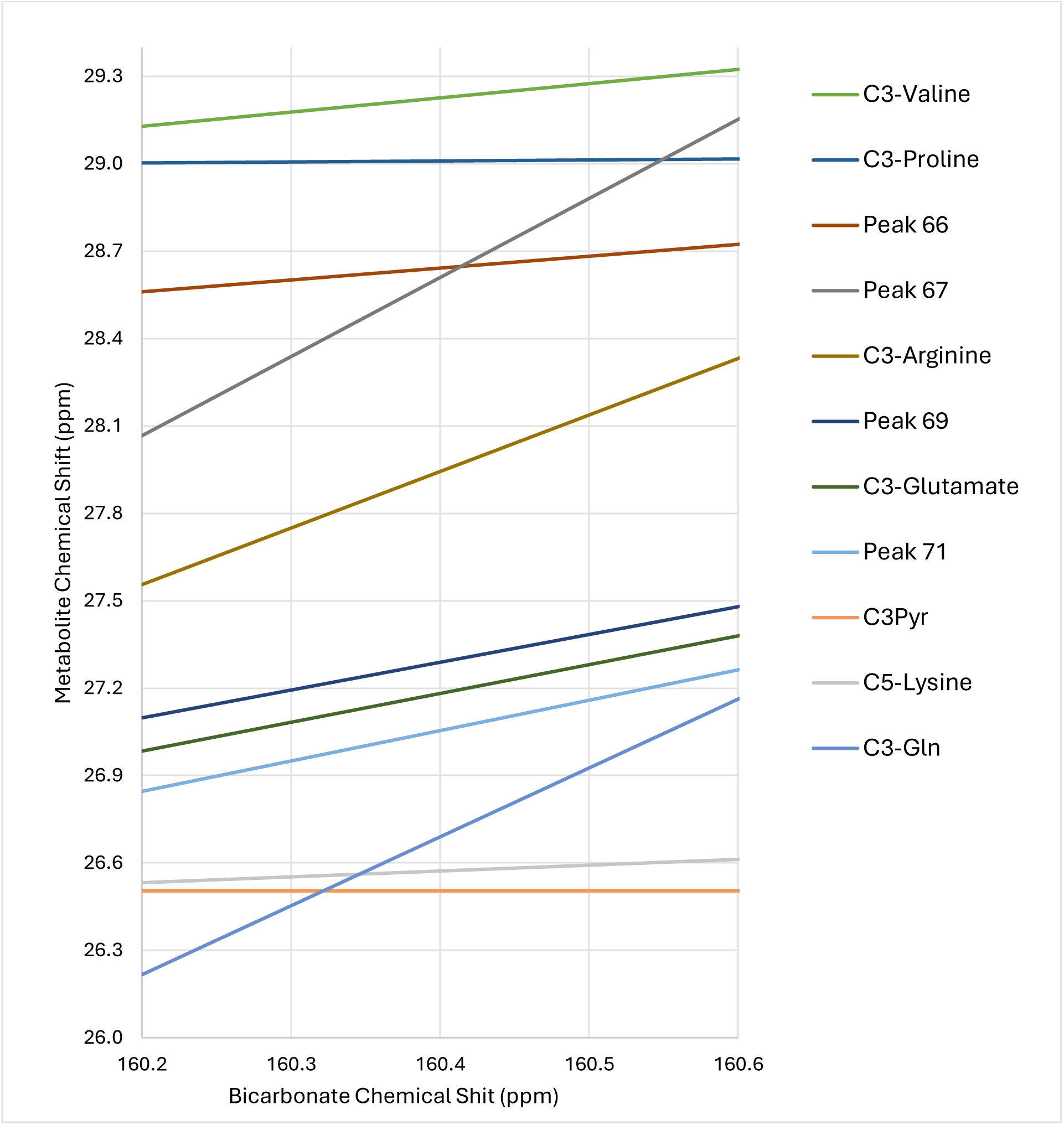
Simulations of Chemical Shift Variance. Simulated ^13^C chemical shifts demonstrate that certain peaks can be unambiguously assigned across the studied pH range due to non-overlapping chemical shifts (e.g., C_3_-valine). Other peaks exhibit overlapping only at specific pH values but remain resolvable when the bicarbonate reference shift is known (e.g., C_3_-arginine, Peak 69, C_3_-glutamate). A subset of peaks, such as Peak 66 (C_3_-glutamine), show partial overlap across the pH range, allowing for unambiguous assignment only under certain conditions.

**Table S2:** Metabolic Flux Ratios and ^13^C Enrichment for all treatment groups.

|  |  | With L-Glutamine |  |  |  |  | Without L-Glutamine |  |  |  |  | All |  |  |  |  |
| --- | --- | --- | --- | --- | --- | --- | --- | --- | --- | --- | --- | --- | --- | --- | --- | --- |
|  |  | WT |  | LIS |  | p value * | WT |  | LIS |  | p value * | WT |  | LIS |  | p value * |
| Flux Ratio:<br>Intermolecular | $^{13}\text{C}_1\text{-Glc}/^{13}\text{C}_3\text{-Lac}$ | 1.46 | ± 0.14 | 1.44 | ± 0.19 | | 1.28 | ± 0.12 | 1.44 | ± 0.30 | | 1.39 | ± 0.15 | 1.44 | ± 0.20 | |
| | $^{13}\text{C}_2\text{-Ac}/^{13}\text{C}_3\text{-Lac}$ | 0.053 | ± 0.033 | 0.072 | ± 0.048 | | 0.052 | ± 0.017 | 0.081 | ± 0.037 | | 0.052 | ± 0.025 | 0.076 | ± 0.040 | |
| | $^{13}\text{C}_3\text{-Ala}/^{13}\text{C}_3\text{-Lac}$ | 0.035 | ± 0.010 | 0.035 | ± 0.008 | | 0.016 | ± 0.001 | 0.015 | ± 0.002 | | 0.027 | ± 0.012 | 0.026 | ± 0.012 | |
| | $^{13}\text{C}_3\text{-Ser}/^{13}\text{C}_3\text{-Lac}$ | 0.020 | ± 0.010 | 0.030 | ± 0.011 | | 0.029 | ± 0.017 | 0.037 | ± 0.028 | | 0.024 | ± 0.013 | 0.033 | ± 0.018 | |
| | $^{13}\text{C}_3\text{-Pyr}/^{13}\text{C}_3\text{-Lac}$ | 0.019 | ± 0.015 | 0.019 | ± 0.021 | | 0.020 | ± 0.0164 | 0.015 | ± 0.008 | | 0.020 | ± 0.0129 | 0.017 | ± 0.016 | |
| | $^{13}\text{C}_3\text{-Form}/^{13}\text{C}_3\text{-Lac}$ | 0.0036 | ± 0.0012 | 0.0033 | ± 0.0012 | | 0.0037 | ± 0.0013 | 0.0042 | ± 0.0016 | | 0.0036 | ± 0.0011 | 0.0037 | ± 0.0014 | |
| | $^{13}\text{C}_4\text{-Glu}/^{13}\text{C}_3\text{-Lac}$ | 0.0074 | ± 0.0021 | 0.0121 | ± 0.0031 | p=0.05 | 0.0043 | ± 0.0010 | 0.0054 | ± 0.0010 | | 0.0061 | ± 0.0023 | 0.0092 | ± 0.0042 | |
| | $^{13}\text{C}_2\text{-Glu}/^{13}\text{C}_3\text{-Lac}$ | 0.0020 | ± 0.0004 | 0.0037 | ± 0.0008 | p=0.009 | 0.0025 | ± 0.0011 | 0.0033 | ± 0.0009 | | 0.0022 | ± 0.0007 | 0.0035 | ± 0.0008 | p=0.006 |
| | $^{13}\text{C}_3\text{-Glu}/^{13}\text{C}_3\text{-Lac}$ | 0.0015 | ± 0.0003 | 0.0024 | ± 0.0004 | p=0.008 | 0.0012 | ± 0.0003 | 0.0015 | ± 0.0006 | | 0.0014 | ± 0.0003 | 0.0020 | ± 0.0007 | p=0.04 |
| Flux Ratio:<br>Intramolecular | $^{13}\text{C}_3/^{13}\text{C}_1\text{-Lac}$ | 61 | ± 4 | 65 | ± 5 | | 65 | ± 5 | 66 | ± 7 | | 64 | ± 6 | 66 | ± 5 | |
| | $^{13}\text{C}_3/^{13}\text{C}_2\{^{12}\text{C}_3\}\text{-Lac}$ | 61 | ± 4 | 66 | ± 4 | | 67 | ± 8 | 80 | ± 7 | | 66 | ± 8 | 72 | ± 9 | |
| | $^{13}\text{C}_3/^{13}\text{C}_2\{^{13}\text{C}_3\}\text{-Lac}$ | 95 | ± 6 | 116 | ± 20 | p=0.10 | 102 | ± 9 | 115 | ± 9 | | 101 | ± 9 | 116 | ± 15 | p=0.05 |
| | $^{13}\text{C}_4/^{13}\text{C}_2\text{-Glu}$ | 3.8 | ± 0.6 | 3.3 | ± 0.1 | | 1.9 | ± 0.2 | 1.7 | ± 0.2 | | 3.0 | ± 1.1 | 2.6 | ± 0.9 | |
| | $^{13}\text{C}_4/^{13}\text{C}_3\text{-Glu}$ | 5.1 | ± 0.9 | 4.9 | ± 0.7 | | 3.5 | ± 0.3 | 4.4 | ± 1.5 | | 4.4 | ± 1.1 | 4.7 | ± 1.0 | |
| | $^{13}\text{C}_2/^{13}\text{C}_3\text{-Glu}$ | 1.4 | ± 0.2 | 1.5 | ± 0.1 | | 2.0 | ± 0.4 | 2.5 | ± 1.0 | | 1.6 | ± 0.4 | 1.9 | ± 0.8 | |
| 13C<br>Enrichment: | $\text{C}_3/\text{C}_2\text{-Ala}$ | 28 | ± 3 | 22 | ± 4 | | 17 | ± 3 | 16 | ± 9 | | 23 | ± 7 | 20 | ± 7 | |
| | $\text{C}_3/\text{C}_2\text{-Ser}$ | 11 | ± 1 | 12 | ± 5 | | 12 | ± 4 | 14 | ± 7 | | 12 | ± 3 | 13 | ± 6 | |
| 13C<br>Enrichment:<br>Absolute | $\text{C}_1\text{-Glc}$ | 0.98 | ± 0.01 | 0.98 | ± 0.01 | | 0.98 | ± 0.01 | 0.98 | ± 0.01 | | 0.98 | ± 0.01 | 0.98 | ± 0.01 | |
| | $\text{C}_3\text{-Lac}$ | 0.42 | ± 0.01 | 0.42 | ± 0.01 | | 0.41 | ± 0.02 | 0.40 | ± 0.04 | | 0.42 | ± 0.01 | 0.41 | ± 0.03 | |
| | $\text{C}_2\text{-Ac}$ | 0.27 | ± 0.10 | 0.25 | ± 0.12 | | 0.31 | ± 0.04 | 0.31 | ± 0.06 | | 0.28 | ± 0.08 | 0.28 | ± 0.09 | |
| | $\text{C}_1\text{-Form}$ | 0.12 | ± 0.02 | 0.11 | ± 0.02 | | 0.15 | ± 0.05 | 0.16 | ± 0.06 | | 0.13 | ± 0.04 | 0.12 | ± 0.04 | |
| *only p<0.1 are shown for simplicity |  |  |  |  |  |  |  |  |  |  |  |  |  |  |  |  |

**Table S3:**
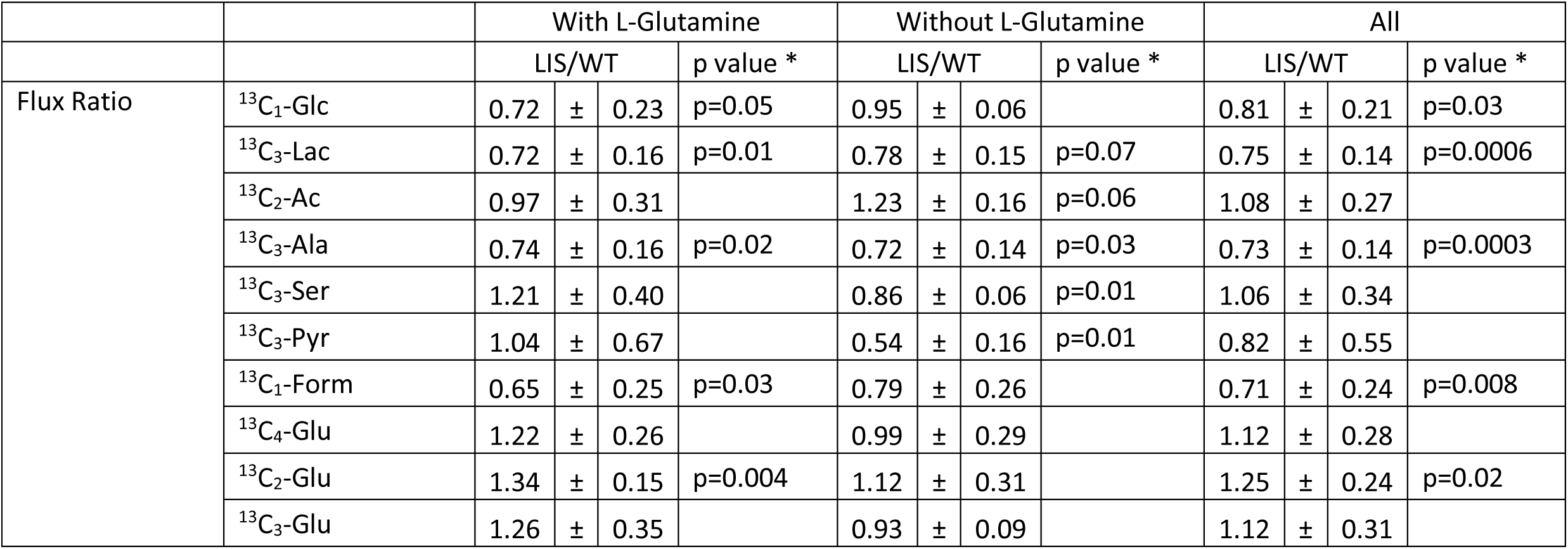
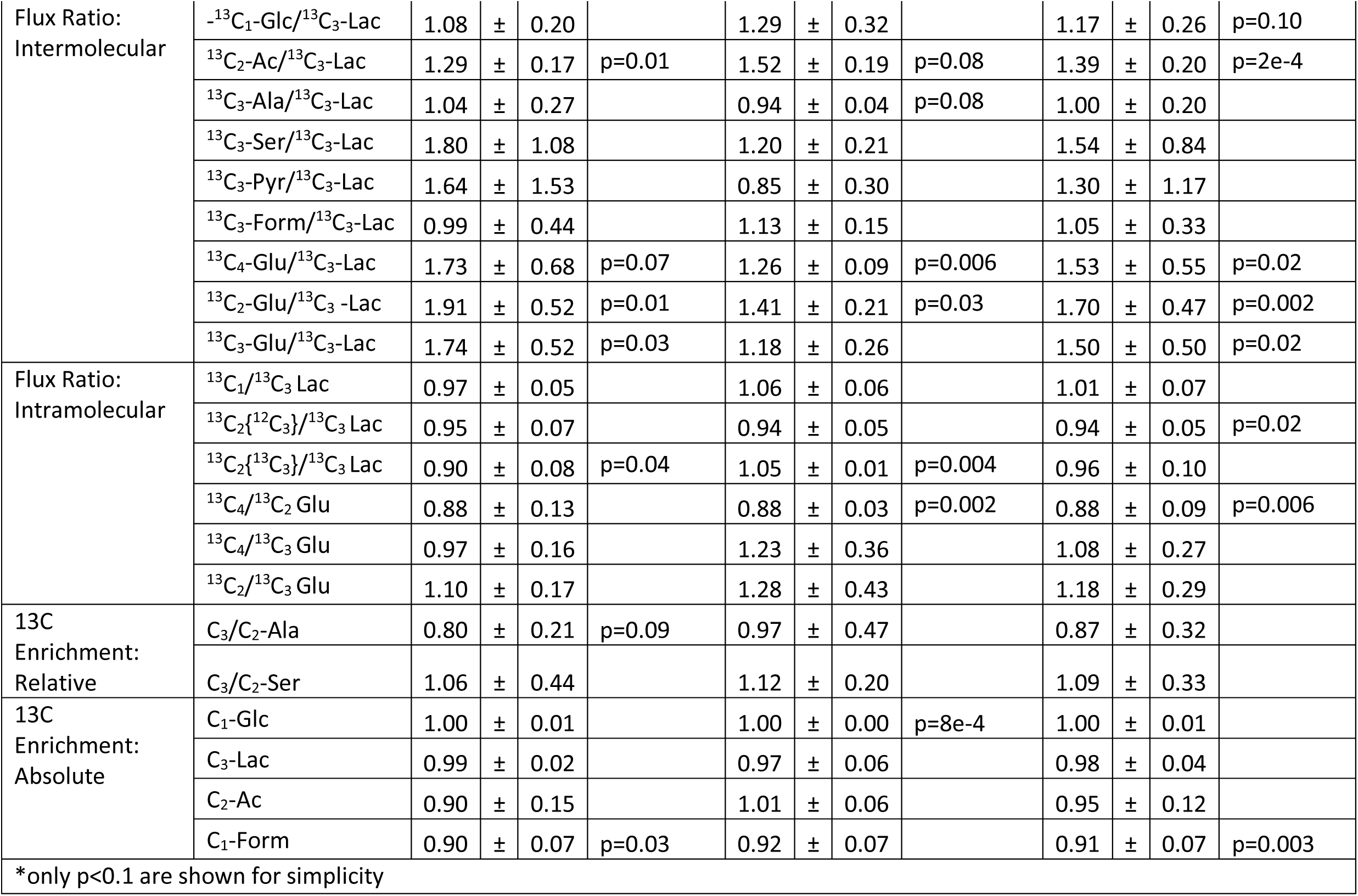
Ratio of Metabolic Fluxes, Metabolic Flux Ratios and ^13^C Enrichment for sets of WT and LIS hESCs cultured in parallel.

**Figure S2:**
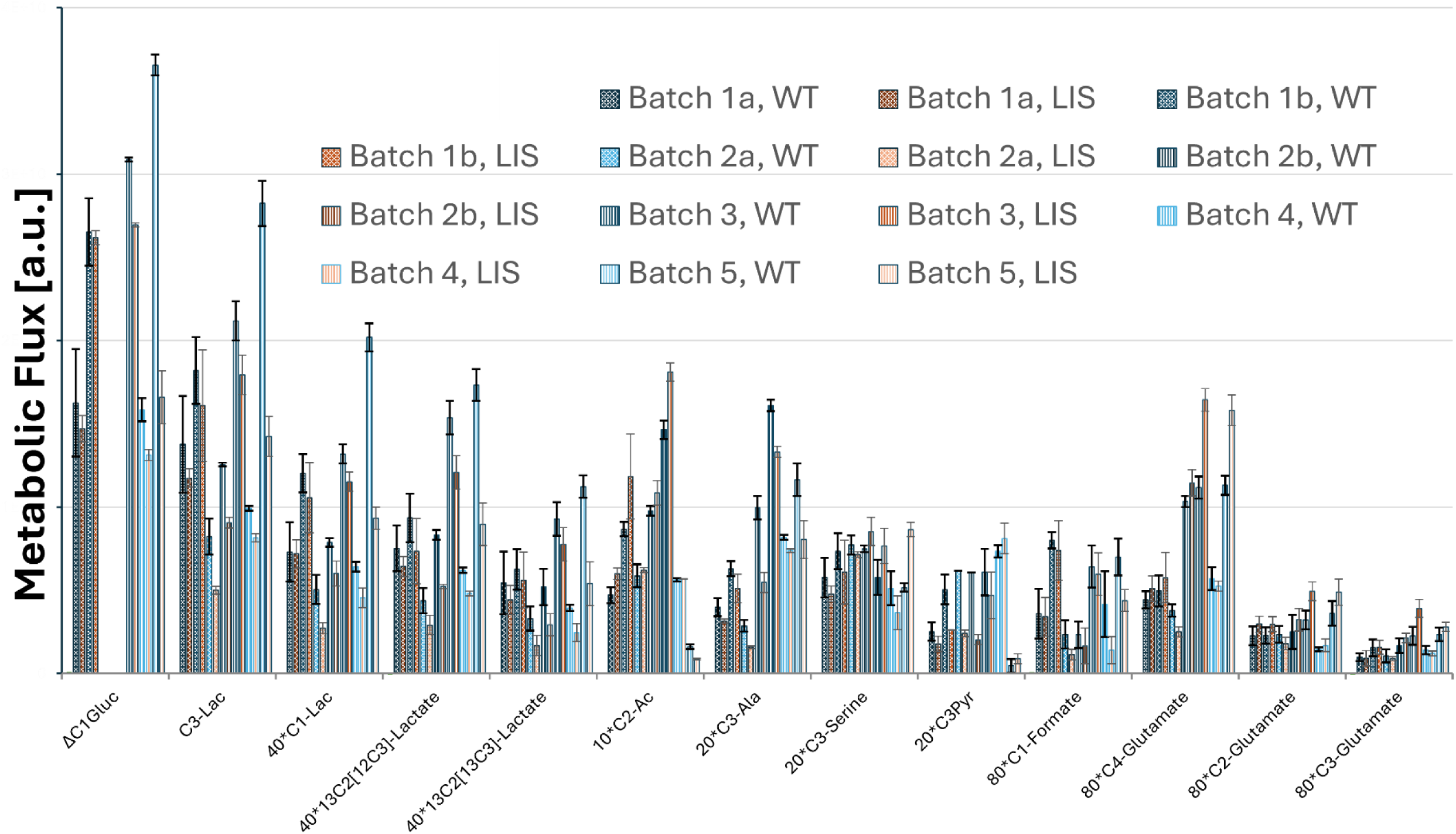
Metabolic Flux for all sets of hESCs.

**Figure S3:**
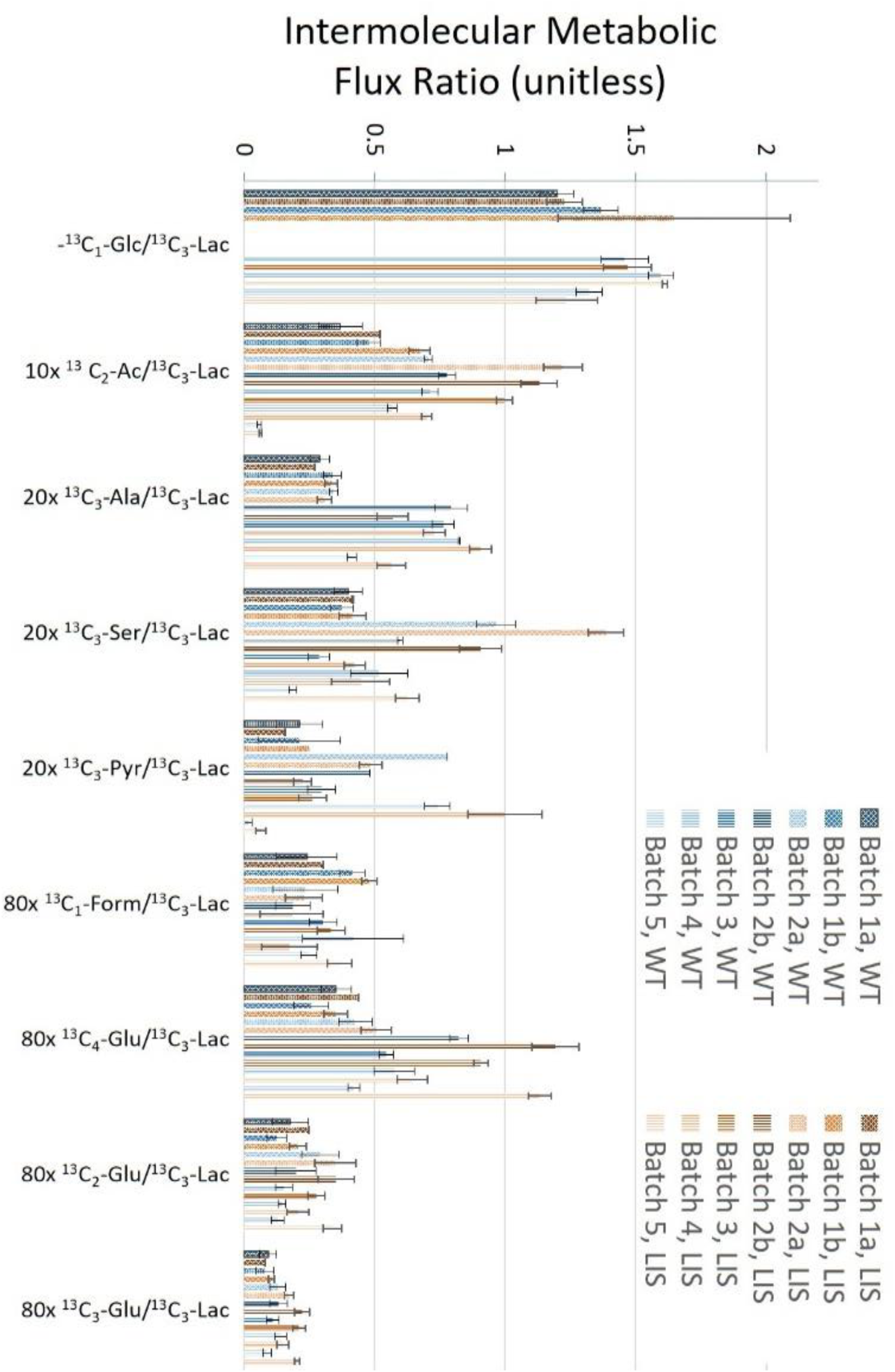
Intermolecular Metabolic Flux Ratios for all sets of hESCs.

**Figure S4:**
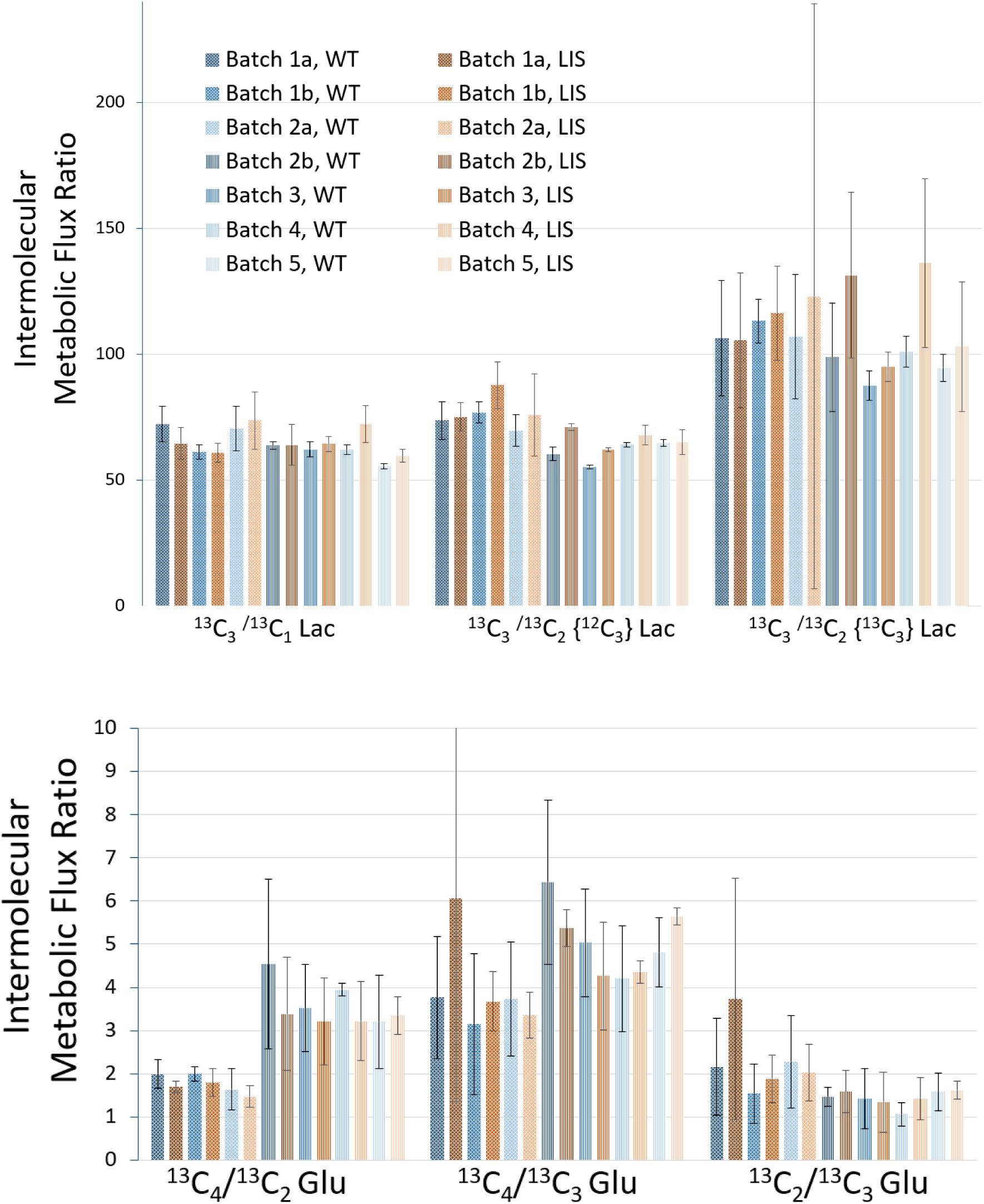
Intramolecular Metabolic Flux Ratios for all sets of hESCs.

**Figure S5:**
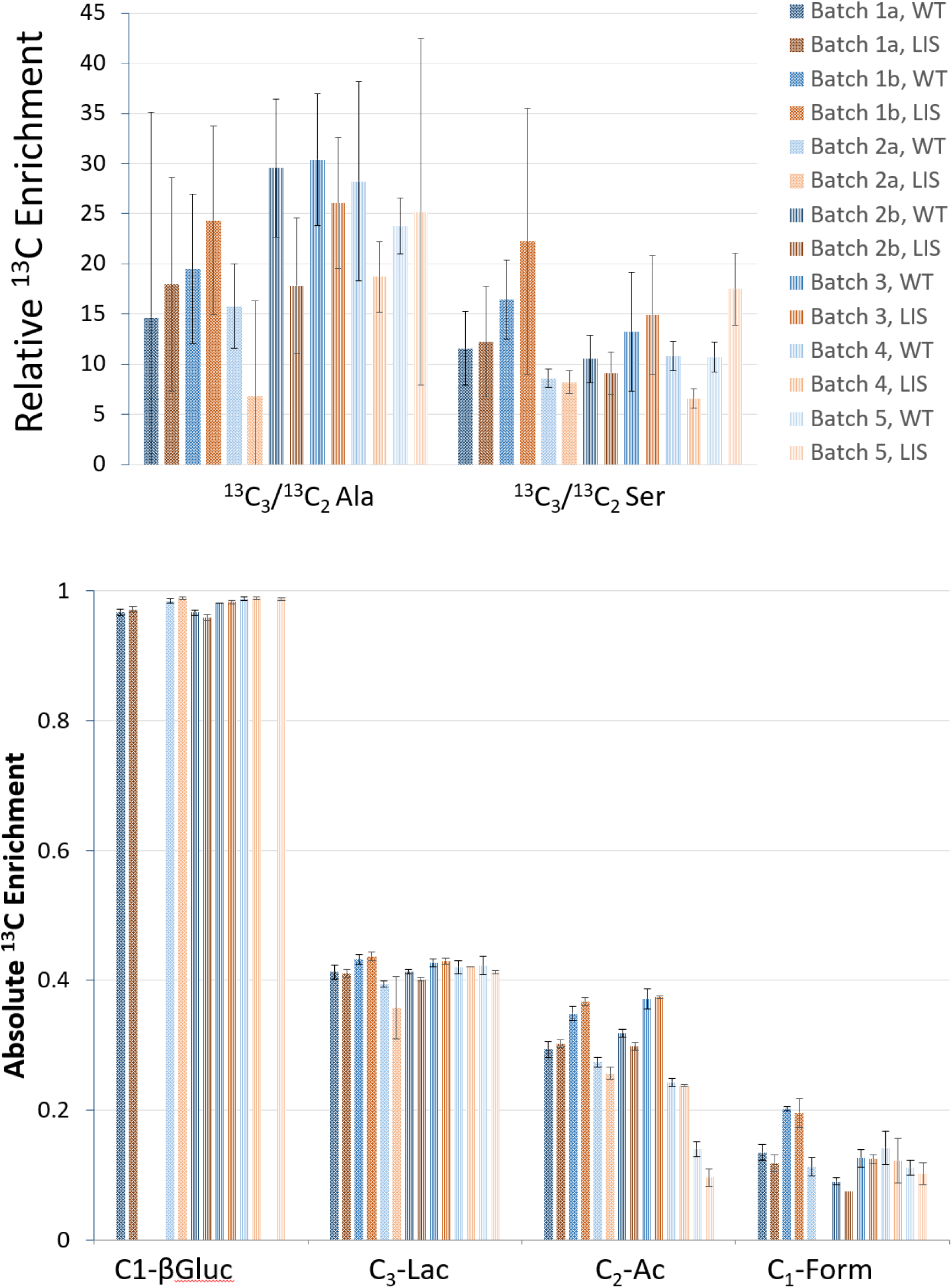
Relative and Absolute 13C enrichment for all sets of hESCs.

